# Multi-omics Profiling of the Lateral Ventricle Choroid Plexus Reveals Developmental Cellular Remodeling, Early Immune Gene Activation, and a Novel Epithelial Subtype

**DOI:** 10.1101/2025.08.25.671514

**Authors:** Mónica Fernandes, Lili Andersson-Li, Eduarda Correia, Alice Vieira, Diana Alves-Afonso, Ana Isabel Álvarez-López, Jonas Campos, Neemat Mahmud, Mandy Meijer, Eneritz Agirre, Fernanda Marques, Claudia Nobrega, Gonçalo Castelo-Branco, João Carlos Sousa, Ana Mendanha Falcão

## Abstract

Healthy brain development and function highly depend on the choroid plexus. Temporal alterations in the cellular landscape and gene expression of choroid plexus cells can alter immune cell trafficking in the brain and cerebrospinal fluid composition, ultimately impacting brain dynamics. Here, we performed a comprehensive multi-omics analysis—including bulk and single-cell transcriptomics and epigenomics—of the lateral ventricle choroid plexus across early postnatal and adult stages in mice and rats. We uncovered striking changes in the choroid plexus cellular composition from neonatal to adult stages, accompanied by transcriptional remodeling of all main cell types. Immune cells were markedly increased in adulthood and immune cell profiling revealed an altered cell-type diversity through time. Surprisingly, we observed an early gene activation of host-defense genes in all choroid plexus main cell types, beginning in the neonatal period and progressively increasing into young adulthood. Moreover, some genes induced in epithelial cells in response to inflammation were found to be epigenetically primed, despite not being transcriptionally active. Epithelial cells exhibited subtype diversity and plasticity, with distinct gene expression programs and chromatin accessibility profiles emerging over time. Notably, we identified a novel epithelial cell subtype with unique gene markers suggesting a specialized function potentially linked to neuro-signaling. Ligand-receptor interaction analysis revealed a progressive remodeling of cellular crosstalk networks during choroid plexus maturation, suggesting dynamic intercellular signaling as the tissue develops.

Our study offers a comprehensive atlas of transcriptional activity and chromatin accessibility in choroid plexus cells, providing a valuable resource to guide future efforts in targeting gene expression at the choroid plexus for therapeutical purposes.

## INTRODUCTION

The choroid plexus (CP) is a unique tissue essential for brain homeostasis, known for producing cerebrospinal fluid (CSF) to support brain buoyancy and forming the blood-CSF barrier through a monolayer of epithelial cells connected by tight junctions. In recent years, an expanding range of functions have been attributed to the CP, broadening its initially perceived role. In health, CPs have been implicated in diverse functions ranging from the regulation of lymphocyte trafficking for immune surveillance (Baruch and Schwartz, 2013; Shipley et al., 2020) to CSF-mediated functions such as modulation of cortical development (Lehtinen et al., 2011), adult neurogenesis (Falcão et al., 2012; Silva-Vargas et al., 2016), brain plasticity (Spatazza et al., 2013), anxiety-like behavior (Vincent et al., 2021), learning and memory (Arnaud et al., 2021) and fine tuning of the circadian clock (Myung et al., 2018; Quintela et al., 2018). CPs are also recognized to remove metabolic and toxic waste from the brain and to act as a signaling center from where messages are both received and transmitted to and from the brain (Kaiser and Bryja, 2020).

To carry out these functions, CPs rely on four major cell types traditionally characterized as epithelial, mesenchymal, vascular and immune cells. Characterizing the heterogeneity of CP cell populations and their compositional changes, transcriptional and chromatin landscape profiles over time is crucial to comprehend how such a small tissue can effectively accommodate a multitude of functions with impact on brain function.

A recent report utilizing single-cell/nucleus RNA-sequencing has significantly advanced our understanding of the cellular diversity within CPs, revealing ventricle-specific and age-related differences across embryonic, adult, and aged stages (Dani et al., 2021). This study uncovered a shared embryonic progenitor that gives rise to both epithelial and neuronal lineages, alongside age-dependent transcriptional programs in epithelial and fibroblast cells—including a pronounced activation of host-defense genes in aged epithelial populations. In the aged CP particularly, it also revealed intricate cell-cell communication networks, such as cell-type-specific IL-1β signaling, pointing to emerging immune crosstalk with aging (Dani *et al*., 2021). Despite these advances, key aspects of CP maturation—particularly during a critical neonatal window—and the chromatin dynamics driving its development and cellular heterogeneity remain largely unexplored.

Here, we present a comprehensive multi-omics analysis of CP maturation during a critical developmental window—from neonatal stages where myelination and synaptogenesis occur, to young adulthood—in two rodent species (*Mus musculus* and *Rattus norvegicus*). Using bulk RNA-seq in rats, single-cell RNA-seq (scRNA-seq), and single-cell multiome (ATAC + gene expression), in mice, we investigated how CP cell composition, heterogeneity, and chromatin accessibility evolve after birth, with a particular focus on the lateral ventricle. Our results reveal remarkable transcriptomic changes across all cell types, accompanied by shifts in chromatin accessibility and major alterations in cell type proportions from early postnatal stages to adulthood. We observe a progressive acquisition of immune-like gene signatures in endothelial, mesenchymal, and epithelial cells already at neonatal stages, in parallel with a remarkable change in immune cell-types diversity over time. Through epigenomic profiling, we further identified inflammation-associated genes in epithelial cells that, while not actively expressed in healthy conditions, were already epigenetically primed for activation— highlighting a poised state for rapid immune response.

Notably, proliferative endothelial and mesenchymal populations were restricted to the neonatal period, suggesting active postnatal remodeling of CP vasculature and stroma. In addition, we identify a marker for epiplexus cells in the tissue and characterize a novel epithelial cell subtype expressing *Gata3* and *Atp10b*, potentially linked to neuronal signaling. Lastly, our ligand-receptor interaction analysis highlights dynamic changes in cell-cell communication within the lateral ventricle CP, suggesting that intercellular signaling is progressively remodeled during CP cell maturation.

Our study reveals a highly dynamic cellular and transcriptional landscape of the lateral ventricle CP during development, which might impact the CP itself and the brain, and provides a valuable resource for future research into CP function and pathology.

## RESULTS

### Multi-omics transcriptomic and chromatin accessibility landscapes of the lateral ventricle CP cells in neonatal and adult ages

To investigate the processes underlying lateral ventricle CP maturation from neonatal stages to adulthood, we have initially conducted a bulk RNA-seq experiment in Wistar Han rats at postnatal day (P) 1, 4, 7, 10 and 60 (Figure 1A). Principal component analysis demonstrated a clear segregation from P1 and P60, with P4, P7 and P10 presenting more similarities and being intermediates between the other stages (Figure 1B). In accordance, differential gene expression analysis showed higher differences in neonatal stages vs P60 (Figure S1A, Table S1). When comparing P1 vs P60, gene ontology analysis on the biological function showed enrichment in P1 for pathways related to extracellular matrix organization, cell division, tissue development while at P60 we found an enrichment in antigen processing and presentation, response to external stimulus, and fatty acid metabolic process pathways (Figure S1B, Table S1). Similarly, P10 vs P60 comparison showed cell differentiation, cell cycle, system development pathways enrichment in P10 while at P60 we find immune response, response to cytokine, regulation of body fluid levels pathways (Figure S1B, Table S1). Interestingly, the upregulation of adaptive immune system starts earlier at postnatal stages, as observed by the increase of adaptive immune genes already at P10 vs P1 (Figure S1B, Table S1).

**Figure 1:**
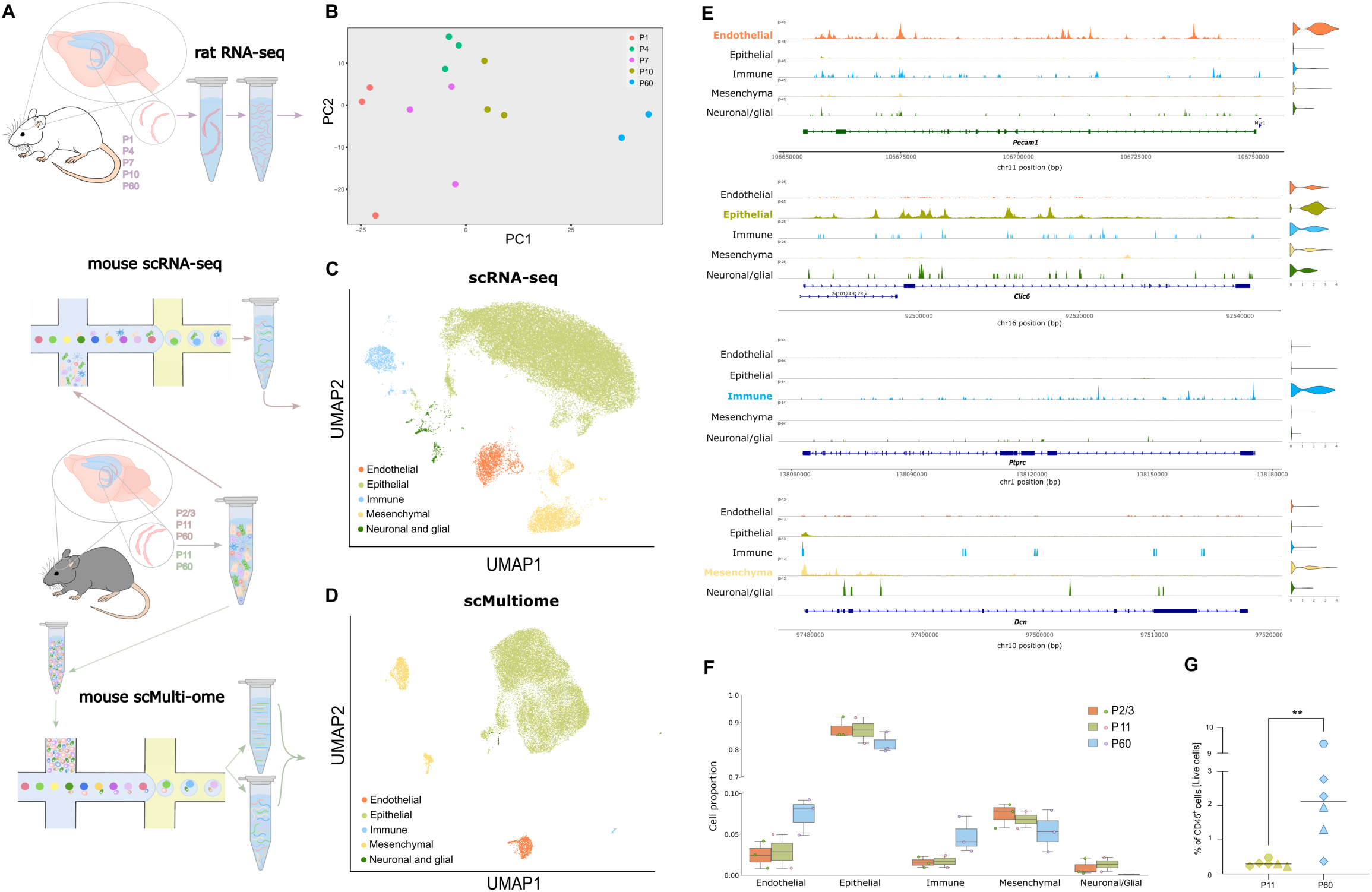
Multi-ome analysis of the lateral ventricle CP from mouse and rat at different ages. (A) Schematic representation of the experimental set up for the multi-ome analysis of mouse and rat lateral ventricle CP at different time points. (B) Principal component analysis (PCA) plot of bulk RNA-seq data from rat lateral ventricle CP across five color-coded time points. Each dot represents the lateral ventricle CP from a single rat. (C) Uniform manifold approximation and projection (UMAP) clustering of scRNA-seq data from mouse lateral ventricle CP cells, color-coded by major cell types and comprising samples from postnatal day (P)2/3 (n=3), P11 (n=2), and P60 (n=3). (D) UMAP clustering of scMultiome (simultaneous scATAC and RNA-seq) data from mouse lateral ventricle CP cells, color-coded by major cell types at P11 (n=3) and P60 (n=3). (E) Integrative Genomics Viewer (IGV) tracks for chromatin accessibility for markers of the cell types identified in scMultiome. Violin plots depicting the expression of the genes, and the corresponding genomic coordinates are shown. Top left depicts the range of the normalized signal. (F) Boxplots depicting the different cell-type proportions across P2/3 (n=3), P11 (n=2) and P60 (n=3). (G) Graph plots obtained from flow cytometry analysis of CD45+ immune cells out of total lateral ventricle CP live cells at P11 and P60. Median is shown, n=6, Mann-Whitney test** p<0.01.

To resolve the cellular identities and associated transcriptome of CP through time, we next performed single-cell RNA-seq (scRNA-seq) in lateral ventricle CP of mice at three selected time points P2-3, P11 and P60 (Figure 1A, S2B). We acquired 47550 lateral ventricle CP cells passing quality control (QC) (Figure S2A) and identified six major cell types according to previously described canonical markers: epithelial (*Folr1*, *Otx2*), mesenchymal (*Lum*, *Dcn*), endothelial (*Plvap*, *Pecam1*), immune (*Fcer1g*, *C1qa*) and neuronal/glial cells (*Sox6*, *Plp1*) (Figure 1C, S2C). To further characterize the epigenetic landscapes of these cells along with the chromatin modifications occurring at postnatal maturation processes, we also performed single-cell Multiome (scMultiome: scRNA-seq in combination with scATAC-seq), of lateral ventricle CP from P11 and P60 mice (Figure 1A, S2E). In this experiment we have also taken advantage of a TTR-eGFP knock-in mice, *Ttr* is a well-recognized marker of CP epithelial cells whose presence in other cell types is perceived as a contamination (Olney et al., 2022) which we have confirmed in our data (Figure S2H-I). After QC (Figure S2D) we successfully mapped the chromatin accessibility and transcriptome of 53835 cells. Transcriptome and differential chromatin accessibility analysis led to the identification of the six main cell types (Figure 1D, S2E-F), in accordance with scRNA-seq. Nevertheless, the amount of immune, neuronal and glial cells captured in the multiomic assay was much lower than in the single cell RNA-Seq approach (Figure 1D).

Chromatin accessibility for cell-types was also determined, and observed mainly in intronic, intergenic and promoter regions for all cell types (Figure S2G). Based on the accessibility of marker genes, we have found increased and/or restricted chromatin accessibility at the cell-type gene markers (Figure S2F), as depicted with IGV tracks for *Clic6* locus (epithelial cells), *Pecam1* (endothelial), *Dcn* (mesenchymal) and *Ptprc* (immune cells) (Figure 1E). As anticipated, the TTR promoter was accessible almost exclusively in epithelial cells and expression of TTR promoter-driven eGFP expression was exclusive from epithelial cells (Figure S2H-I).

### Cell type composition of the lateral ventricle CP is altered from neonatal to adult ages

While cellular proportions were very similar between P2/3 and P11, at P60 we observed an increase in the proportion of endothelial and immune cells. These changes were balanced by a gradual decrease in epithelial and mesenchymal cells proportions in lateral ventricle CP (Figure 1F).

To confirm the increase of the immune cell population in adult stages we performed flow cytometry analysis of lateral ventricle CP cells from six P11 and six P60 mice and observed a sharp increase in the CD45^+^ immune cells in adult stages, in agreement with our scRNA-seq data (Figure 1G). Interestingly, gene ontology analysis from rat bulk RNA-seq showed enrichment in genes related to B cell chemotaxis and leukocyte chemotaxis at P60 when comparing with P1 and P10, respectively (Figure S1B), which could be explained by the observed increase in immune cells in adult ages. Altogether, these findings suggest that lateral ventricle CP maturation is a dynamic and conserved process across rodent species.

### Early activation of host-defense genes in all CP cell types

To investigate whether the early upregulation of host-defense genes observed in rat bulk RNA-seq was likely due to an increased number of immune cells, as suggested by the scRNA-seq data, activation of these genes within CP resident cells, or a combination of both, we analyzed the expression patterns of adaptive and innate immune genes across individual CP cell types over time. We found upregulation of immune genes that was cell-type-dependent. For instance, MHC class I genes gradually increased in mesenchymal and endothelial cells, while MHC class II genes showed an overall increase across the populations, though with varying expression levels. The chemokines *Cxcl12* and *Cx3cl1* were upregulated in mesenchymal and endothelial cells, while interferon-related genes *Ifi27* and *Ifi27l2a* increased in epithelial, immune, and mesenchymal cells (Figure 2A). Despite the marked upregulation of MHC class I and II gene expression from P11 to P60, chromatin accessibility at their respective loci in P60 cells showed minimal or no corresponding increase (Figure 2B).

**Figure 2:**
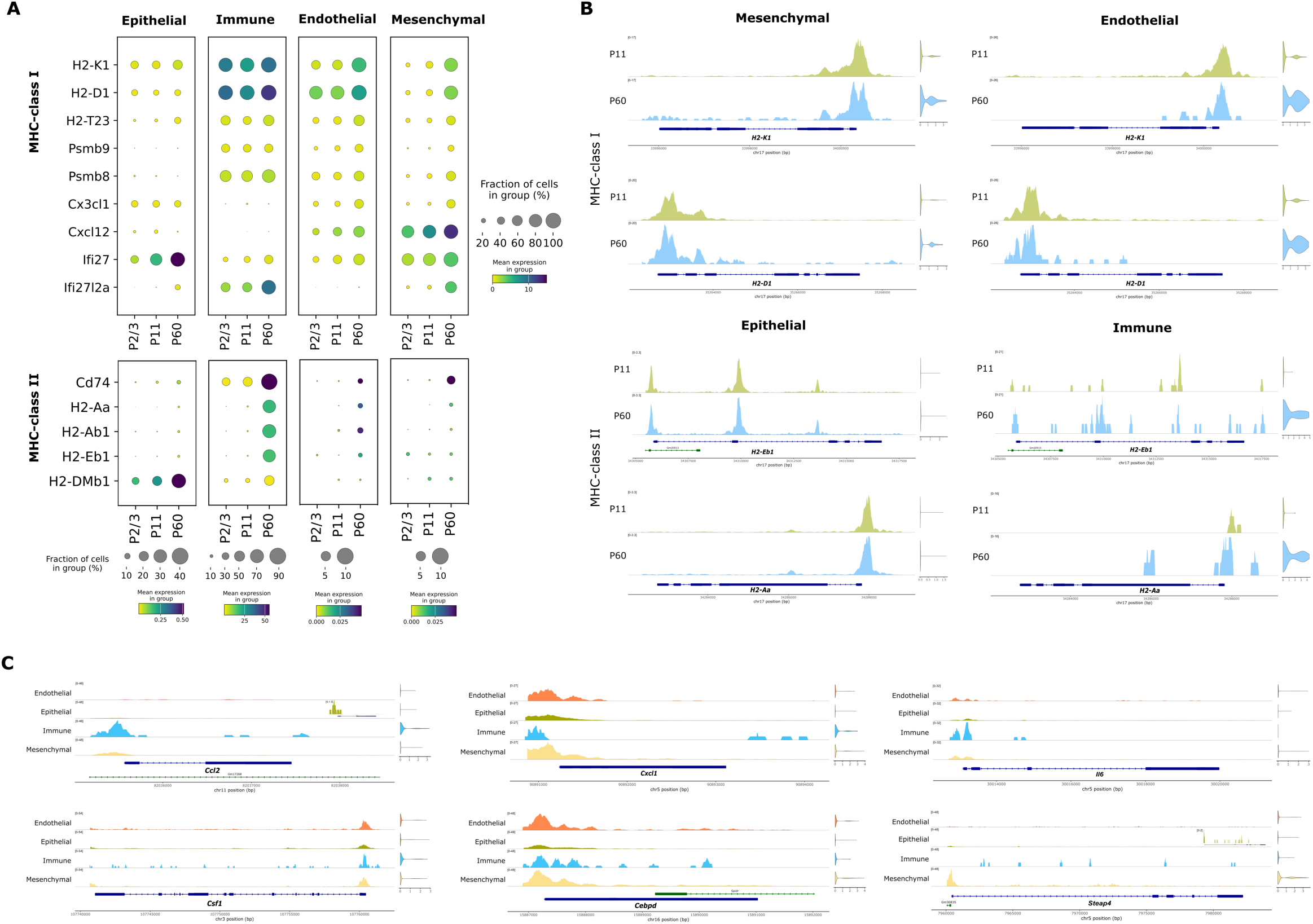
Early immune gene activation in lateral ventricle CP cells and epigenetic priming of inflammation-elicited genes in epithelial cells. (A) Dot plot of MHC class I and II, and immune-related genes across time and cell populations at the lateral ventricle CP. (B) IGV tracks for chromatin accessibility of across cell populations at P11 and P60 time points. Violin plots depicting the expression of the genes, and the corresponding genomic coordinates are shown. (C) IGV tracks for chromatin accessibility of genes in lateral ventricle CP cell populations. Represented genes were previouly reported to be upregulated in response to acute inflammation in epithelial cells. Violin plots depicting the expression of the genes, and the corresponding genomic coordinates are shown.

Our results align with previously reported age-associated immune signatures in the CP (Baruch et al., 2014; Dani *et al*., 2021; Mesquita et al., 2015), although our data shows that such changes begin as early as P10/11.

### Epigenetic Priming of Inflammation-Responsive Genes in Epithelial Cells

Previous studies have shown that CP epithelial cells transiently and quickly alter their transcriptome in response to inflammation (Marques et al., 2009a; Xu et al., 2024). This process is considered essential for the CP’s adaptation to inflammation and is thought to support immune cells by regulating their trafficking across the blood-CSF barrier and modulating immune responses (Xu *et al*., 2024). Both peripheral and intraventricular administration of lipopolysaccharide has triggered the expression of immune response genes in epithelial cells, which are virtually absent from their basal transcriptome under healthy conditions (Marques *et al*., 2009a; Xu *et al*., 2024). Here, we profiled the chromatin accessibility of a subset of these genes in the lateral ventricle CP cells and found that *Csf1*, *Cxcl1* and *Cebpd* exhibited open chromatin across CP cell types but had very limited or no expression under healthy conditions (Figure 2C). In contrast, *Steap4* displayed open chromatin in mesenchymal cells, while showing reduced chromatin accessibility in other cell types. Similarly, chromatin accessibility at the *Ccl2* and *Il6* loci was predominantly restricted to immune cells. Notwithstanding, a subset of epithelial cells also exhibited chromatin accessibility at the promoter regions of these genes (Figure 2C), which explains their rapid and transient induction upon lipopolysaccharide stimulation. It remains an open question whether lipopolysaccharide-mediated inflammation induces increased chromatin accessibility in additional epithelial cells or if only those with pre-existing open chromatin respond, potentially serving as a mechanism to prevent a cytokine storm.

### Immune cell profile of the lateral ventricle CP remarkably changes in adult mice

Our analysis of the vascular, epithelial and mesenchymal cells of the lateral ventricle CP indicates that these cells gradually transition to an immune-prone profile already at neonatal stages to adulthood. We thus examined the immune cell compartment at the lateral ventricle CP, in scRNA-seq of three time points. We identified nine distinct populations: dendritic cells (DC) (expressing high levels of MHC-II genes such as *H2-Dmb1*), neutrophils (*S100a9*, *S100a8*), classical monocytes (*Emilin2, Chil3*), non-classical monocytes (*Adgre4*, *Gfpt1*), B cells (*Cd79a*, *Igkc*), NK cells (*Gzmb, Klrb1c*), T cells (*Cd3d*, *Lat*), macrophages (*Ctsb*, *Mrc1*) and epiplexus cells (*Syngr1, Crybb1*) (Figures 3A and B, S3A and B). The presence of these immune cell types agrees with recent reports on CP immune cell populations (Dani *et al*., 2021; Van Hove et al., 2019).

**Figure 3:**
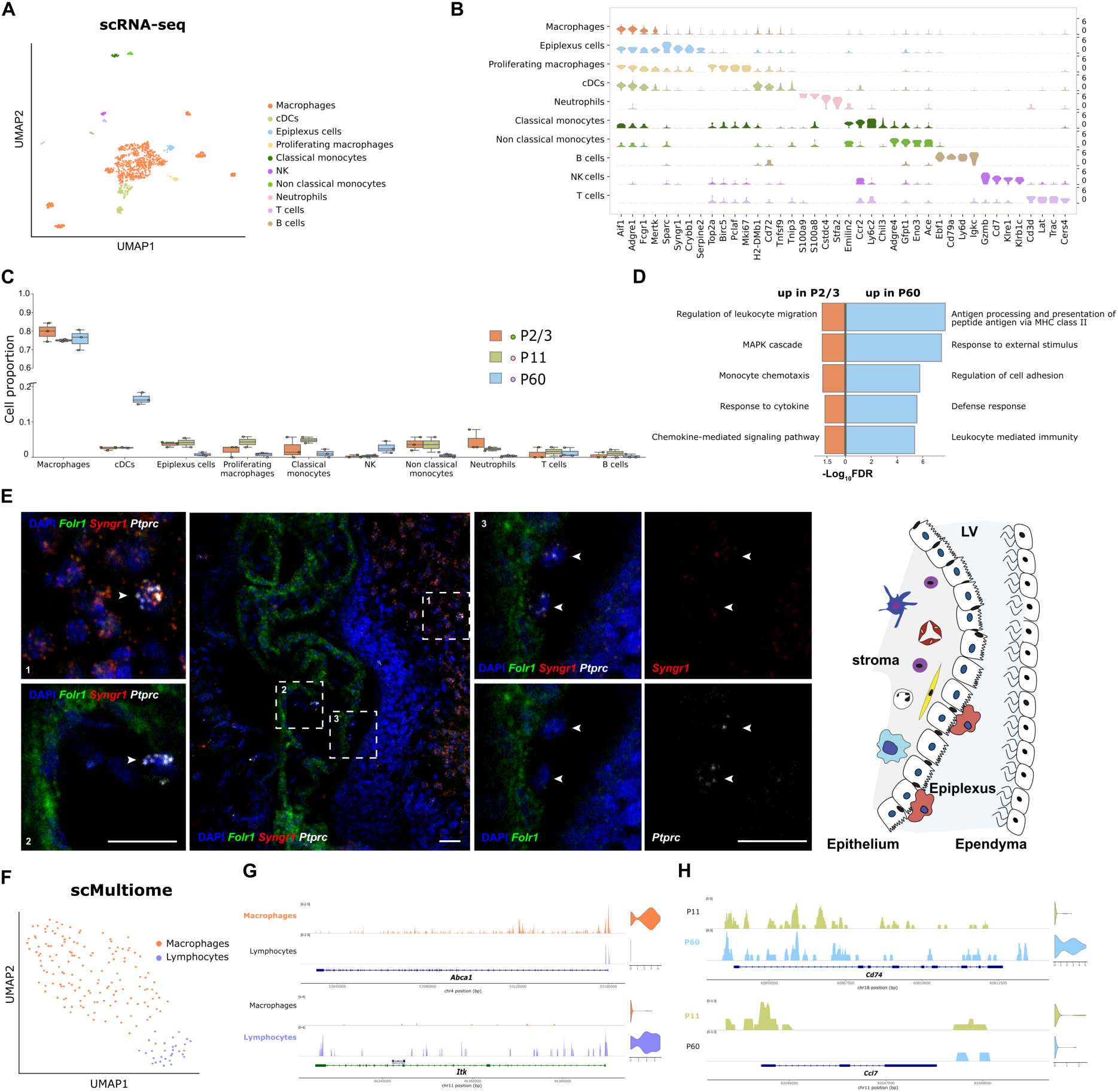
Immune cell-type diversity within lateral ventricle CP and fluctuations across different ages. (A) UMAP of immune cells color-coded by cell-type, and comprising three time points, P2/3 (n=3), P11 (n=2) and P60 (n=3). cDCs, classic Dendritic Cells. (B) Violin plots representing the median expression levels of four markers for the lateral ventricle CP immune cell-types. (C) Boxplots representing immune cell-type proportions across three time points, P2/3 (n=3), P11 (n=2) and P60 (n=3). (D) Gene ontology analysis (GO) for the biological function of genes enriched in P2/3 and P60 stages of mouse lateral ventricle CP macrophages. (E) RNAscope in situ hybridization (ISH) representing a tissue section of lateral ventricle CP marked with probes for *Fo/r1* (epithelial marker), *Ptprc* (immune cell marker) and *Syngr1* (epiplexus cell marker). Dashed boxes shown at higher magnification highlight regions of *Syngr1+/Ptpcr+* cells in brain parenchyma (microglia), *Syngr1-/Ptpcr+* in lateral ventricle CP stroma (stromal immune cells) and *Syngr1+/Ptpcr+* lying over the epithelial apical side (epiplexus cells). Representative images, n = 3 biologically independent P11 mouse lateral ventricle CP. Scale bars, 20 µm. (F) UMAP clustering of immune lateral ventricle CP cells, from scMultiome (simultaneous scATAC and RNA-seq), color-coded by cell-type, and comprising two time points, P11 (n=3) and P60 (n=3). (G) IGV tracks for chromatin accessibility for markers of the macrophages and lymphocytes identified in scMultiome. Violin plots depicting the expression of the genes, and the corresponding genomic coordinates are shown. (H) IGV tracks for chromatin accessibility of differentially expressed genes in P11 and P60, in macrophages identified in scMultiome. Violin plots depicting the expression of the genes, and the corresponding genomic coordinates are shown.

We identified a transcriptional signature for epiplexus cells—CSF-contacting macrophage-like cells—that allows their reliable identification without the need for spatial localization within the CP tissue. Epiplexus cells expressed *Syngr, Crybb1, Sparc*, markers also reported to be expressed in microglial cells. We have validated by RNAscope in situ hybridization (ISH), *Syngr1* gene as a marker that distinguishes epiplexus cells from other macrophages from lateral ventricle CP (Figure 3E and S3C). Of note, previous reported data of scRNA-seq of mouse brain macrophages suggested that epiplexus cells expressed all microglial signatures genes but at lower levels (Van Hove *et al*., 2019). In accordance, we have observed *Syngr1* molecules in microglia (*Ptprc^+^*) from brain parenchyma (Figure 3E). Surprisingly, we also detected a sporadic *Syngr1^+^* cell bridging the brain parenchyma and the lateral ventricle CP (Figure S3C), which raises the hypothesis that these cells can move directly between these compartments and might even originate from microglia itself instead of shared ontogeny as suggested before (Van Hove *et al*., 2019).

We then estimated the proportions of each immune cell type in neonatal and adult lateral ventricle CP and found striking fluctuations of cell types with age. While neutrophils, non-classical monocytes and epiplexus cells were enriched in neonatal time points, NK cells were present in adult lateral ventricle CP but nearly absent in neonatal mice (Figure 3C). Moreover, the proportion of dendritic cells increased approximately 3 times in adults while non-classical macrophages decreased. B cells, T cells, classical monocytes and proliferating macrophages did not change their proportions from neonatal to adult ages (Figure 3C). The increase in dendritic cells and NK cells in CP in adult stage coincides with the increased expression of immune genes in epithelial, vascular and mesenchymal cells, which could suggest a role of the immune cells in the observed transitions to immune-like states.

Differential expression analysis for macrophage population revealed significant differences at the transcriptome of neonatal and adult lateral ventricle CP (Figure S3D and Table S2). Again, the transcriptome changes were gradual over age. Interestingly, gene ontology analysis shows that at neonatal stages macrophages are enriched in genes involved in monocyte chemotaxis, regulation of leukocyte migration suggesting that at this stage the lateral ventricle CP immune niche is dynamic (Figure 3D). At adult stage macrophages upregulated pathways of defense response and antigen processing and presentation, which were also reported before when comparing embryonic and adult CP (Dani *et al*., 2021), suggesting that upregulation of this pathway persists beyond embryonic stages and occurs gradually over time.

Profiling immune cells of lateral ventricle CP with scMultiome was challenging as we have acquired very few cells in comparison with scRNA-seq (Figure 1C and D). This could be due to a loss of these cells during the procedure and because we had significantly lower amounts of adult lateral ventricle CP cells which are enriched in immune cells (Figure 1F and G, S2E). Nevertheless, we could identify macrophage and lymphocyte lineage cells at the transcriptome (Figures 3F and S3E-G) and at the chromatin level (Figures 3G). When comparing chromatin accessibility in the macrophages of genes up and downregulated, we found that although gene expression is significantly increased in some genes at adult stages, our data suggests that the chromatin accessibility is not altered at P11, which could suggest epigenetic priming. Conversely, *Ccl7* exhibits enriched chromatin accessibility at P11, in agreement with transcriptomic data (Figure S3D).

### Endothelial and mesenchymal cell type composition and gene expression profiles change from neonatal to adult stages in the lateral ventricle CP

We next analysed endothelial and mesenchymal cells across the different ages, P2/3, P11 and P60 (Figure 4A and S4A-C) and divided the endothelial cell population in capillary cells (*Plvap*, *Esm1*), veins (*Vwf*, *Pcdh7*) and arteries (*Pcsk5*, *Nebl*), and mesenchymal cells in pericytes (*Kcnj8*, *Abcc9*), vascular smooth muscle cells (VSMC) (*Acta2*, *Tagln*), fibroblasts (*Col26a1*, *Tcf21*) and leptomeningeal fibroblasts (*Slc6a13*, *Clec3b*) (Figure 4A and B). Importantly, we identified proliferating endothelial cells (*Top2a*, *Mki67*) that gradually decreased over time and were nearly absent in adult stages. Accordingly, the proliferating fibroblasts were present in neonatal and undetected in adults (Figure 4C). Our results suggest active remodeling of the lateral ventricle CP stroma and active angiogenesis at neonatal stages. The presence of a proliferative endothelial population is in line with the previously reported at embryonic CP (Dani *et al*., 2021) and further supports that angiogenesis remains active at least until P11 stages. In the mesenchymal cell populations, while pericytes and leptomeningeal fibroblasts were decreased, VSMC cells proportions remained relatively stable through time (Figure 4C).

**Figure 4:**
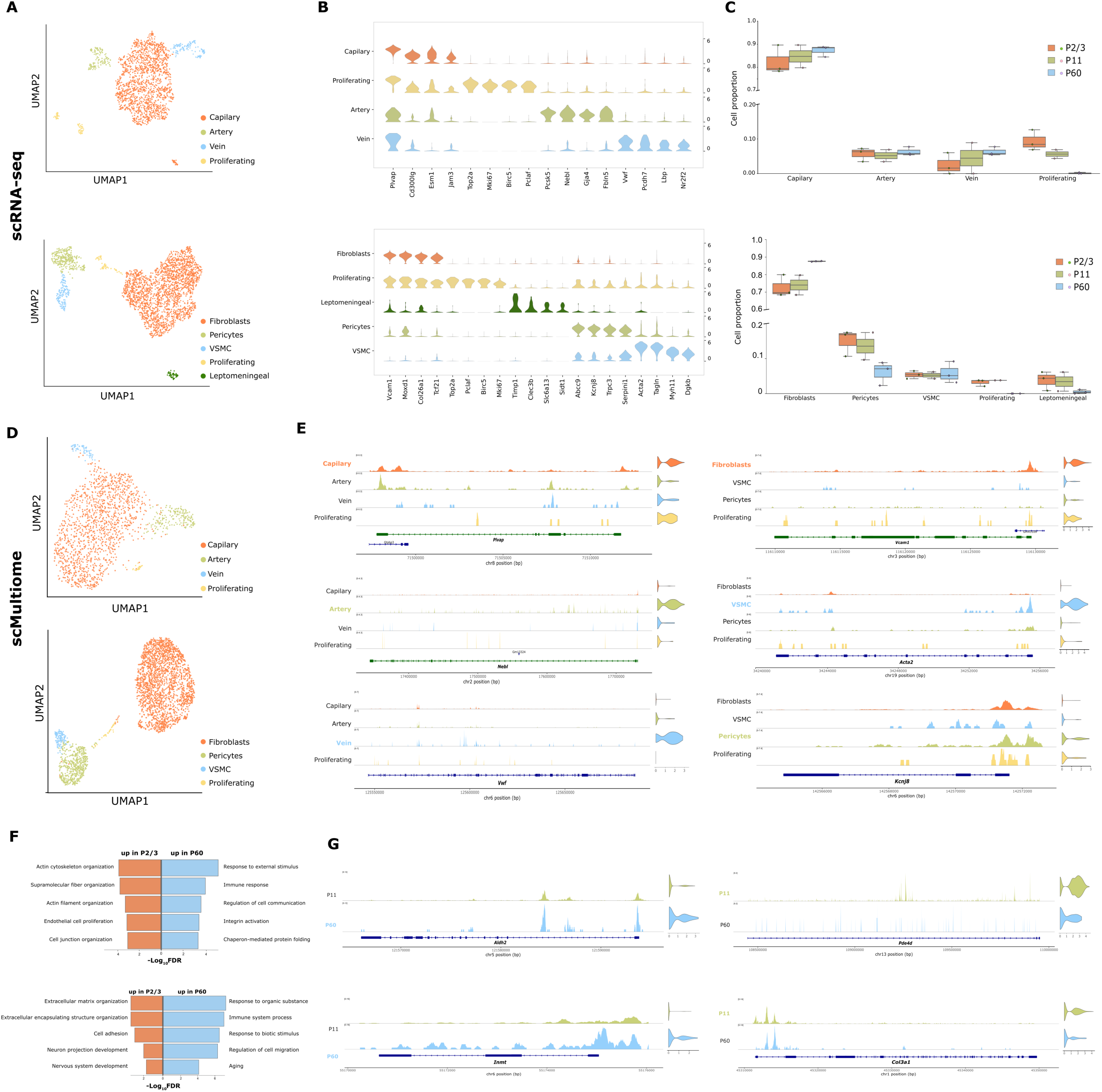
Temporal dynamics of the transcriptome and chromatin accessibility in distinct endothelial and mesenchymal cell types of the lateral ventricle choroid plexus. (A) UMAP of endothelial and mesenchymal cells from scRNĀseq, color-coded by cell-type, comprising three time points, P2/3 (n=3), P11 (n=2) and P60 (n=3). (B) Violin plots representing the median expression levels of four selected markers in each endothelial and mesenchymal cell type. (C) Boxplots representing endothelial and mesenchymal cell-type proportions across three time points, P2/3 (n=3), P11 (n=2) and P60 (n=3). (D) UMAP clustering of endothelial and mesenchymal cells, from scMultiome (simultaneous scATAC and RNA-seq), color-coded by cell-type, comprising two time points, P11 (n=3) and P60 (n=3). (E) IGV tracks for chromatin accessibility of the endothelial and mesenchymal cell-type markers identified in scMultiome. Violin plots on the right depict the expression of the genes, the corresponding genomic coordinates are shown. (F) Gene Ontology analysis of biological functions associated with genes enriched at the P2/3 and P60 stages in mouse endothelial cells (top) and fibroblast cells (bottom). (G) IGV tracks for chromatin accessibility of differentially expressed genes in P11 and P60, in endothelial cells (top) and mesenchymal cells (bottom) identified in scMultiome. Violin plots depicting the expression of the genes, and the corresponding genomic coordinates are shown.

Analysis of scMultiome at P11 and P60 revealed the endothelial and mesenchymal cell-types in the lateral ventricle CP (Figures 4D and S4C-E) and further discern at the chromatin level (Figures 4E and S4G). As such, arteries exhibited enriched chromatin accessibility at *Nebl* and veins at *Vwf* locus. Capillaries markers displayed similar chromatin accessibility (Figure 4E), which was expected given that these genes are present in all endothelial cells but enriched in this population (Figure 4B). We were able to resolve at the chromatin level the different mesenchymal cells, as observed by enriched chromatin accessibility of marker genes as follows: *Vcam1*, *Acta2* and *Kcjn8* for fibroblasts, VSMC and pericytes, respectively (Figure 4E). The presence of leptomeningeal fibroblasts in the lateral ventricle CP stroma detected in scRNAseq was not captured in the scMultiome, which could be due to differences in the procedure, or alternatively, cells were captured at the border of meninges, although we could not identified them as the recently discovered CP base barrier fibroblasts (Verhaege et al., 2024).

To investigate transcriptome profile changes in endothelial and fibroblasts cells during lateral ventricle CP maturation, we have performed differential expression analysis and demonstrate a transient change in gene expression along age (Figure S4G, Table S3 and S4). Gene ontology analysis for genes enriched in adult endothelial cells suggest involvement in immune response, integrin activation while genes upregulated at neonatal stages are related to cell junction organization, actin filament organization (Figure 4F). For fibroblast cells, gene ontology analysis revealed that P2/3 stages gene signatures are related to extracellular matrix organization, nervous system development, while at P60 upregulated genes are involved in immune system process, response to organic substance (Figure 4F).

Similarly, some of the differentially expressed genes in endothelial and fibroblast cell populations were also altered in bulk RNA-seq data from rat when comparing P1 vs P60 (Figures S2J and S4H-I), indicating that part of the maturation process is conserved in these two species. Altogether, our data suggests an active remodeling of the stromal vasculature and matrix in the CP during early post-natal stages.

Importantly, some changes at the transcriptome level over time were mimicked at the chromatin level, as observed by the enriched chromatin accessibility of *Aldh2* and *Pde4d* in P60 and P11, respectively, for endothelial cells, and *Inmt* and *Col3a1* in P60 and P11, respectively, for fibroblast cells (Figure 4G). Our data suggests that chromatin landscape changes with age alongside the transcriptome.

### Neurons, but not oligodendrocytes lineage cells, are present in the lateral ventricle CP stroma of mice and rat

We have characterized neuronal and glial populations of the lateral ventricle CP and identified several cell types based on gene expression markers (Figure S5A and B): neurons (*Tubb3, Syt1*), neuroblast-like habenula (*Spink4*, *Tph1*, classified according to (Zeisel et al., 2018), ependymal cells (subdivided in two distinct cluster, expressing either *Ccdc153* and *Wls* or *Krt15* and *Acta2*, based on the previous annotation (Shah et al., 2018)), astrocyte-like cells (*Aldoc, Lrig1*), committed oligodendrocyte precursors (COPs) (*Itpr2*, *Gpr17*), oligodendrocyte precursor cells (OPC) (*Pdgfra*, *Olig2*) and oligodendrocytes (*Mbp*, *Plp1*). The identification of neuronal and glial cell types is in accordance with previously published data on single-cell/nuclei RNA-seq (Dani *et al*., 2021) where embryonic, adult and aged CP from three ventricles were analysed. To confirm the presence of neurons in lateral ventricle CP in both mice and rats, we performed immunofluorescence for ß3-tubulin and neurofilament (NF) and found ß3-tubulin^+^ and NF^+^ cells in both mice and rats (Figure S5C-E). Interestingly, neurons were very abundant in rats and exhibited several axonal varicosities/boutons as observed by both neuronal markers (Figure S5D and E) suggesting the presence of several sites where synapses can occur.

Regarding oligodendrocyte lineage cells we collected the lateral ventricle CP from adult and P11 Sox10:Cre-RCE:LoxP mice, and did not find GFP^+^ cells, indicating that oligodendrocyte lineage cells are absent in the lateral ventricle CP and therefore the observed cells in the scRNA-seq were likely pulled from brain parenchyma together with the CP. Of note, myelin canonical markers, such as *Mbp*, *Mobp*, amongst others, were increased both at P10 and P60 in rat bulk RNA-seq (Figure S1A) and gene ontology analysis shows enrichment in pathways of myelination (Figure S1A and B). To further confirm if these cells were present in rat we have performed CNP staining and observed CNP^+^ cells in the adult rat lateral ventricle CP with a very immature morphology (Figure S5E), that were very similar to isotypic controls but more abundant. We then performed RNAscope ISH for *Sox10* and *Plp1* and found oligodendrocyte lineage cells in rat lateral ventricle CP, although we were unable to confirm their location within the stroma. As such, the detection of these cells in lateral ventricle CP might be the result of dragging from lateral ventricle CP contacts with parenchymal tissue (Figure S5F).

### Epithelial cells exhibit subtype diversity and plasticity along with age-dependent remodeling of cell structure, gene expression, and chromatin accessibility

Epithelial cells are the most abundant cell type in the lateral ventricle CP and have been widely assigned to various functions. Nonetheless, whether specific tasks are accomplished by distinct epithelial subtypes remains unknown. Studies in mice reveal that epithelial cells exhibit a heterogeneous transcriptome across ventricles (Dani *et al*., 2021; Lun et al., 2015) and report a subset of newly differentiated epithelial cells undergoing ciliogenesis, present at embryonic stages (Dani *et al*., 2021). Accordingly, we have observed these cells expressing several genes involved in ciliogenesis (Figure 5A, B, Table S5) and validated by RNAscope ISH with the marker gene *Ccno* in *Folr1^+^*epithelial cells (Figure 5C), herein named ciliogenic epithelial cells. These cells were present in neonatal stages and nearly absent in adults, indicating that newly differentiated epithelial cells continue to be added to the lateral ventricle CP in neonatal stages, a process that is rare in adults (Figures 5D and S6B). Moreover, we have detected a very small subset of proliferating epithelial cells (*Mki67*, *Birc5*), that did not express the ciliogenesis markers (Figure 5A, B and S6C, Table S5). Interestingly, our data uncovered additional subtypes of epithelial cells that were not previously described in rodents (Figure 5A and B). One subtype observed in scRNA-seq was characterized by the expression of *Cp* and *Gpx3*. We performed RNAscope ISH and did not find double *Cp/Gpx3^+^* cells, but instead, we observed a small subset of cells *Gpx3^+^Cp*^-^ dispersed through the lateral ventricle CP that could represent sporadic *Gpx3* expressing cells in the main epithelial cluster (Figure S6D, E). As such, we did not validate this cell subtype in the tissue and hypothesize that they arise in response to tissue processing (in response to damage or mechanical stresses). Despite exhibiting a significant lower number of genes per cell and genes involved in p53 pathway (Figure S6A, E), we did not consider this cluster a low cell viability one, as these cells also displayed lower mitochondrial genes, which is the opposite of what is associated with poor cell viability (Figure S6A).

**Figure 5:**
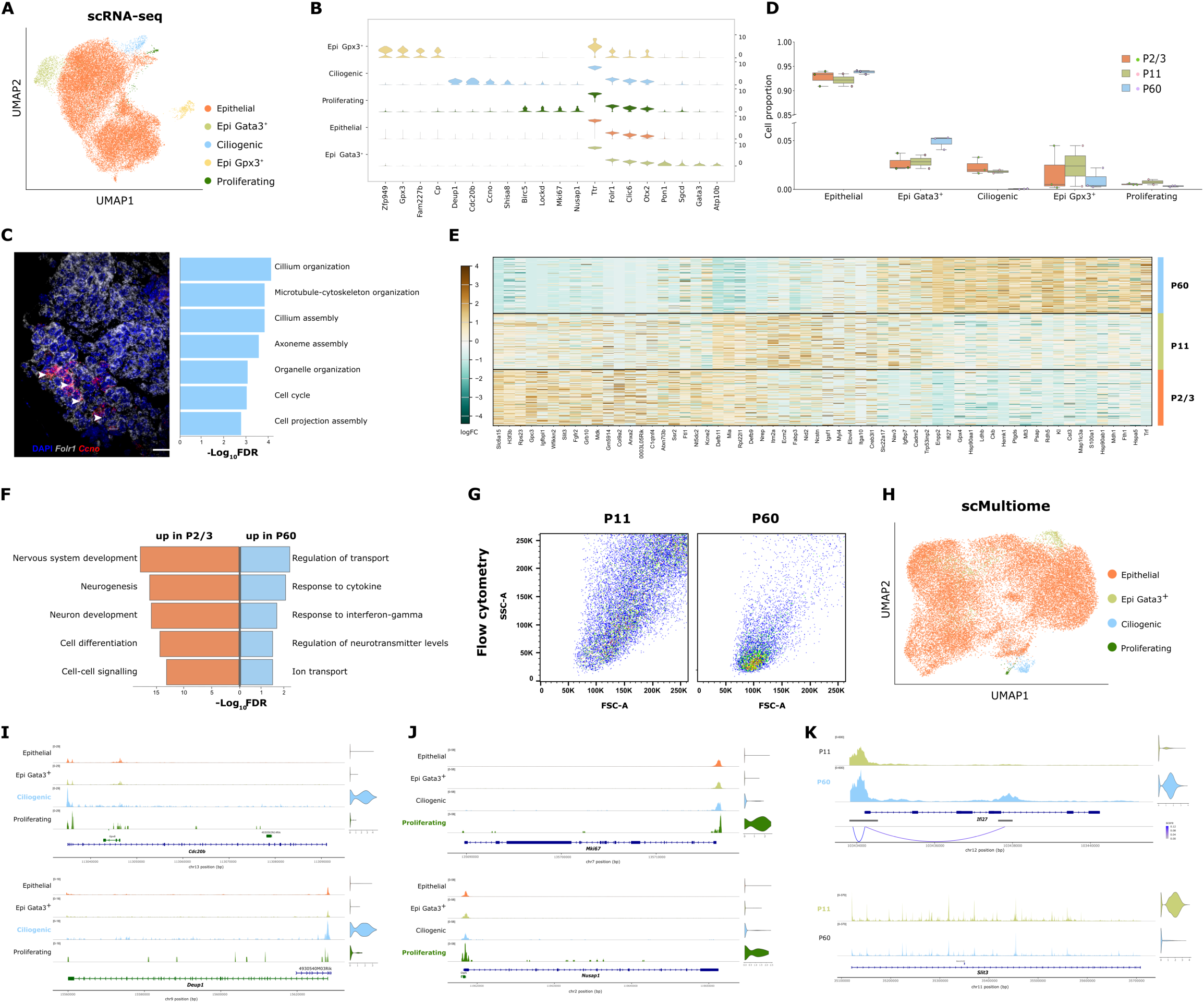
Epithelial cells of the lateral ventricle CP are heterogeneous and exhibit changes in their proportions, transcriptomic profiles, and structural complexity from postnatal to adult stages. (A) UMAP of epithelial cells from scRNA-seq color-coded by cell subtype, comprising three time points, P2/3 (n=3), P11 (n=2) and P60 (n=3). (B) Violin plots representing the median expression levels of four markers for the epithelial cell subtypes. (C) RNAscope ISH for *Folr1+Ccno+* ciliogenic cells in whole mounts from of P11 lateral ventricle CP and GO analysis for the biological function on the top 50 genes enriched in this subtype. Representative images, n = 3 biologically independent P11 mouse lateral ventricle CP. Scale bars, 20 µm (D) Boxplots representing epithelial cell subtype proportions across three time points, P2/3 (n=3), P11 (n=2) and P60 (n=3). (E) Heatmap of differential expression genes in epithelial across different ages. (F) GO analysis for the biological function of genes enriched in P2/3 and P60 stages of mouse epithelial cells. (G) Flow cytometry plots representing cell complexity and size at two different time points, P11 and P60. (H) UMAP clustering of epithelial lateral ventricle CP cell subtypes, from scMultiome (simultaneous scATAC and RNA-seq), color-coded by cell-type, comprising two time points, P11 (n=3) and P60 (n=3). (I) IGV tracks for chromatin accessibility for markers of the ciliogenic cells identified in scMultiome. Violin plots depicting the expression of the genes, and the corresponding genomic coordinates are shown. (J) IGV tracks for chromatin accessibility for markers of proliferating epithelial cells identified in scMultiome. Violin plots depicting the expression of the genes, and the corresponding genomic coordinates are shown. (K) IGV tracks for chromatin accessibility of differentially expressed genes in P11 and P60, in epithelial cells identified in scMultiome. Violin plots depicting the expression of the genes, and the corresponding genomic coordinates are shown.

Differential gene expression analysis on epithelial cells demonstrates a gradual change in transcriptome from neonatal time points to adults (Figure 5E, Table S6). P60 epithelial cells upregulate genes involved in regulation of neurotransmitter levels, regulation of transport and response to cytokine, while earlier time points display genes related to nervous system development, neurogenesis and cell differentiation (Figure 5F). Importantly, variations in gene expression among epithelial cells were partially mirrored by bulk RNA-seq analysis conducted on rats at comparable time points (Figure S6H). Strikingly, flow cytometry analysis of mouse lateral ventricle CP cells reveals a significant difference in the internal complexity and size of epithelial cells (the vast majority of CP cells) from P11 and P60 (Figure 5G). This aligns with the first description of morphological modifications occurring at different stages of choroid plexus maturation in humans, including variations in shape, glycogen granule levels, and other features (Shuangshoti and Netsky, 1966; 1970).

scMultiome identified the same epithelial cell subtypes except for the *Gpx3*^+^/*Cp*^+^ cells, whose existence was not confirmed. Epithelial cell subtypes were segregated in the joint and RNA UMAPs, but not in ATAC-seq UMAP (Figures 5H and S6I-J), suggesting that epithelial cell chromatin landscapes are very similar. To comprehend how differences in transcription are mirrored at the chromatin level we plotted the chromatin accessibility for the epithelial cell subtypes described above. Genes related to ciliogenesis, represented by *Deup1* and *Cdc20b*, exhibited enriched chromatin accessibility in ciliogenic epithelial cells, although their expression was restricted to this specific cell subtype (Figure 5I). A similar pattern was observed in proliferative cell subtypes where the proliferative marker *Mki67* and *Nusap1* were confined to proliferating cells but displayed an opened chromatin in all cell subtypes (Figure 5J).

Next, we investigated whether chromatin accessibility was altered alongside the transcriptome across the two different ages and observed that *Ifi27*, which is increased in young adults, exhibits enriched chromatin accessibility at regulatory regions, while *Slit3* is increased at P11 in agreement with its elevated expression at this stage (Figure 5K).

Our findings suggest that epithelial cells present significant plasticity at the chromatin accessibility level, changing over time, and exhibiting potential to initiate a ciliogenic program and to re-enter the cell cycle upon maturation.

### A new epithelial cell subtype revealed at the lateral ventricle CP

A previously unidentified subtype of epithelial cells was identified by both scRNA-seq and scMultiome experiments. This cell subtype expressed *Gata3, Atp10b, Sgcd*, amongst other gene markers (Figure 5 A, B, S6I-K, Table S4). Accordingly, we found *Gata3*/*Atp10b/Folr1^+^* cells in whole mount lateral ventricle CP (Figure 6A). This subtype comprised about 2.1 to 5.2% of total epithelial cells, depending on the age, as it was increased from neonatal to adult stages (Figure 5D).

**Figure 6:**
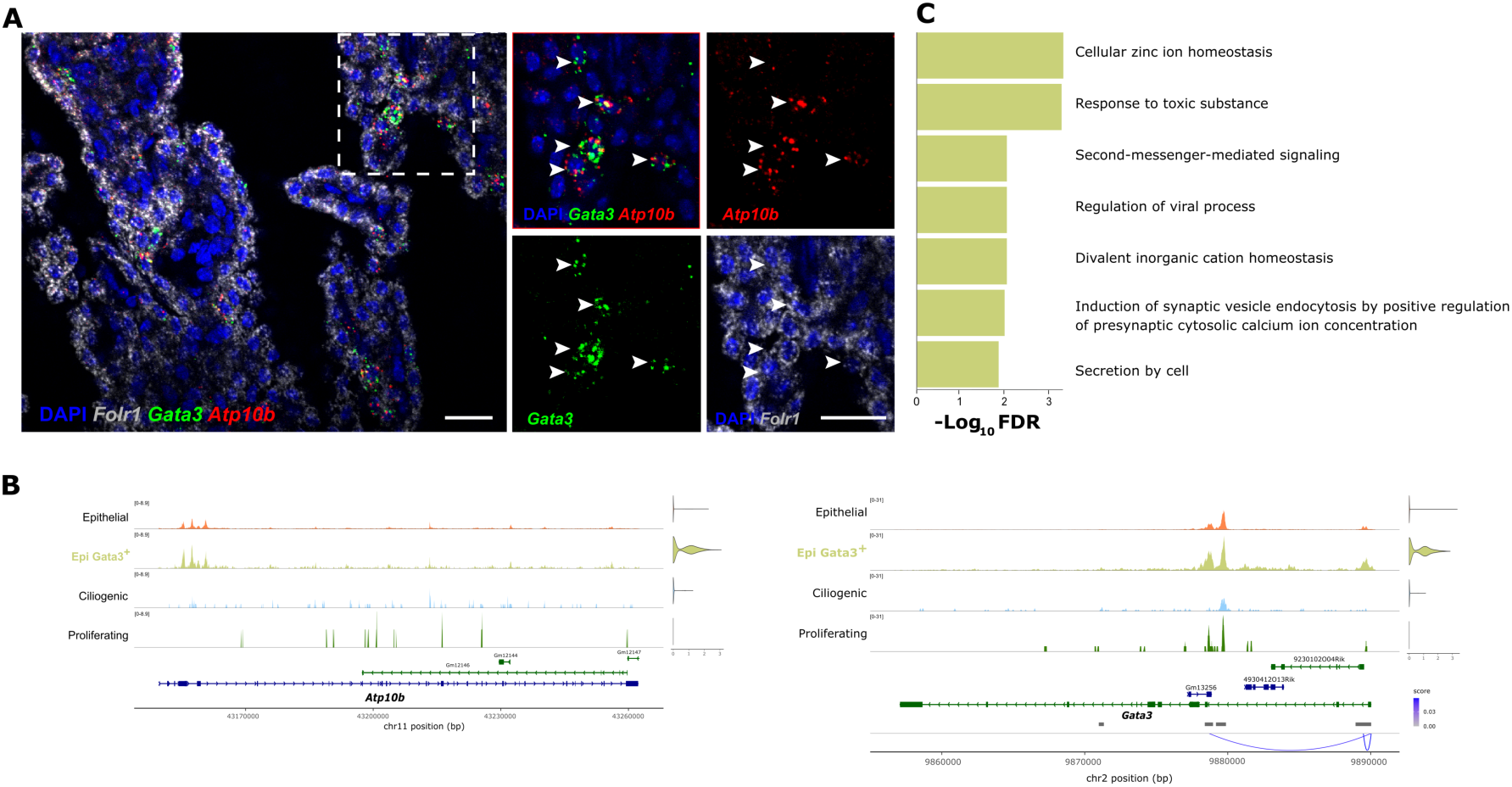
Novel epithelial cell subtype uncovered by scRNĀseq and scMultiomics. (A) RNAscope ISH for *Folr1^+^Atp10b^+^Gata3^+^* cells (arowheads) in whole mounts from of adult lateral ventricle CP. Representative images, n = 3 biologically independent adult mouse lateral ventricle CP. Scale bars, 20 µm. (B) IGV tracks for chromatin accessibility for markers of the Gata3^+^ epithelial cell subtype identified in scMultiome. Violin plots depicting the expression of the genes, and the corresponding genomic coordinates are shown. (C) GO analysis for the biological function on the top 50 genes enriched in this subtype.

Chromatin accessibility was increased at promoter-transcription start site regions of *Gata3* locus in this cell subtype (Figure 6B). While the peaks profile of ciliogenic and proliferating cells were identical across epithelial cell subtypes, *Gata3*^+^epithelial cells displayed a distinct profile compared to ciliogenic cells, as indicated by chromatin accessibility at *Gata3* and *Atp10b* locus (Figure 6B), suggesting that this subtype represents a mature cell state.

We then performed gene ontology analysis on the Biological Function of the top 50 markers and found these cells were enriched in genes related to regulation of viral process, second-messenger mediated signaling, divalent inorganic cation homeostasis, secretion by cell, amongst others (Figure 6C). Importantly, gene ontology analysis on Cellular Component shows that genes enriched in this population are involved in synapse, vesicle membrane, endosome, plasma membrane region, neuron projection (Figure S6G). While the specific function of these cells remains unknown, we hypothesize that they can display a neuronal input-mediated cell secretion that can then be propagated to the neighbor epithelial cells through gap junctions, and/or can represent a cell stage of neuronal activation for apocrine secretion. Interestingly, GATA3 is a transcription factor that in the breast cells is essential for glandular development, driving luminal differentiation and maintaining the identity of mammary luminal epithelial cells (Kouros-Mehr et al., 2006). The presence of Gata3 in the lateral ventricle CP epithelial cells may serve a similar role in maintaining epithelial cell differentiation as they lose part of their cytoplasm during apocrine secretion.

### Ligand-receptor analysis suggests a remodeling of cellular crosstalk in the lateral ventricle during CP cell maturation

Having observed time-dependent changes in gene expression across all CP cell types, we next explored whether cellular crosstalk is also remodeled during maturation. To this end, we applied the Ligand-Receptor Analysis Framework (LIANA) (Dimitrov et al., 2022) to identify potential ligand-receptor interactions among CP cells at P2/3 and P11, aiming to predict how these interaction networks evolve over time. Our analysis revealed that mesenchymal cells emerged as the predominant sender cell type, exhibiting the highest total number of predicted interactions. In contrast, endothelial, mesenchymal, and immune cells were the main receiver populations (Figure 7A). While the overall interaction frequency maps remained relatively stable between P2/3 and P60 (Figure 7A), we visualized the top 40 interactions for the major cell population as sources of ligands, to highlight possible shifts in ligand-receptor communication across time (Figure S7A). Finally, we examined the expression dynamics of these ligand-receptor pairs, focusing on those that showed temporal changes in either the sender or receiver populations (Figure 7B–E).

**Figure 7:**
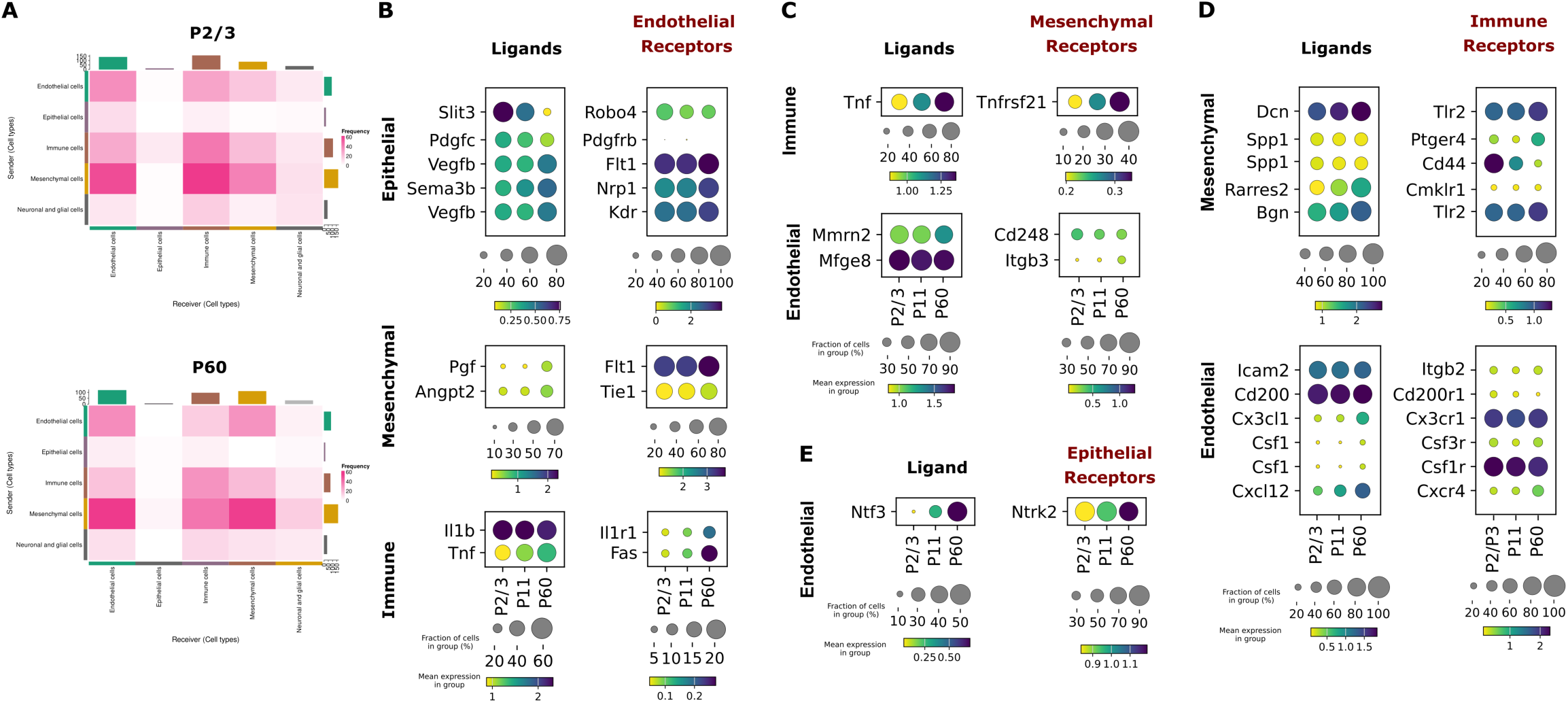
Cell-cell communication in the lateral ventricle CP cells from neonatal to adult stages. (A) Frequency heatmap for the inferred interactions of sender and receiver cell-types, estimated by LIANA. Frequency of interaction (color coded) and sum of interactions (barplots) are depicted. (B) Selected ligands from epithelial, mesenchymal and immune cells to endothelial receptors. Proportion of expressing cells (circle size) and median expression (color coded) is represented for each panel. (C) Selected ligands from immune and endothelial cells to mesenchymal cell receptors. Proportion of expressing cells (circle size) and median expression (color coded) is represented for each panel. (D) Selected ligands from mesenchymal and endothelial cellsto immune cell receptors. Proportion of expressing cells (circle size) and median expression (color coded) is represented for each panel. (E) Selected ligand from endothelial cells to epithelial receptor. Proportion of expressing cells (circle size) and median expression (color coded) is represented for each panel. Ligand receptor pair interactions where selected based on either altered proportion of cells or gene expression magnitude in sender or receiving cell types.

We observed several ligand-receptor pairs involved in angiogenesis, targeting endothelial cells, to be dynamically regulated over time. Notably, the Slit3–Robo4 interaction—previously shown to stimulate endothelial cell proliferation, motility, chemotaxis, and vascular network formation (Dou et al., 2018; Paul et al., 2013; Zhang et al., 2009)—was found to be increased in neonatal stages, gradually reducing with age (Figure 7B), which agrees with our findings of neonatal active angiogenesis. Interestingly, *Il1r1* expression in endothelial cells gradually increased with age (Figure 7B), despite stable *Il1b* levels in immune cells across time. This suggests that, in addition to the previously reported age-associated IL-1β–IL1R1 signaling (Dani *et al*., 2021), there is an early activation of host-defense pathways, as shown in Figure 2. Similarly, the *Tnfrsf21* receptor in mesenchymal cells increased over time, in parallel with its ligand *Tnf* expressed by immune cells (Figure 7C), further supporting the enhanced host-defense signaling during CP maturation and aging (Figure 2).

Likewise, changes in immune signaling pathways were also observed between immune cell receptors and ligands derived from endothelial and mesenchymal cells (Figure 7D). We identified opposing temporal expression patterns for two immune receptors— *Cd44* and *Ptger4*—interacting with the same ligand, *Spp1*, in mesenchymal cells, suggesting different functions. Endothelial-derived ligands such as *Cx3cl1*, *Csf1*, and *Cxcl12* showed increasing expression over time, with their corresponding immune receptors either remaining stable or also increasing (Figure 7D). Notably, both *Cxcl12* and its receptor *Cxcr4*, essential for immune cell chemotaxis, were upregulated in endothelial and immune cells respectively, correlating with the rise in immune cell numbers from neonatal to young adult stages (Figure 1F–G).

Epithelial, neuronal, and glial cells exhibited the fewest predicted interactions. However, we observed a striking increase in the expression of endothelial-derived *Ntf3* and its receptor *Ntrk2* in epithelial cells, suggesting a robust and potentially novel signaling axis whose function in the CP remains to be determined (Figure 7E).

## DISCUSSION

As the CP emerges as both a player and a potential therapeutic target in several neurological disorders, comprehending the physiological maturation processes and the regulatory networks that govern transcriptional programs in CP cells is of utmost importance. Here, we provide a comprehensive analysis of lateral ventricle CP transcriptome and chromatin accessibility in neonatal and adult rodents, unveiling fluctuations in cell type composition, gene expression profiles, and chromatin accessibility over time.

We show the presence of scarce neurons, in mouse lateral ventricle CP and neuronal networks with axonal varicosities in the rat lateral ventricle CP, suggesting an increased complexity in this species. Innervation of CP was first reported in the 19th century (Benedikt, 1874), and since then, the presence of different cholinergic, adrenergic and seretonergic nerves have been described in several species (Dani *et al*., 2021; Lindvall and Owman, 1981). Importantly, whether these nerves are myelinated or not is still poorly discussed. It was suggested that most fibers are unmyelinated, although the presence of Schwann cells on the cat’s lateral ventricle CP was demonstrated using electron microscopy (Edvinsson et al., 1975). We did not confirm the presence of oligodendrocytes as part of lateral ventricle CP stroma cells in mice and rats, although we do not rule out their presence in other mammals.

Lateral ventricle CP innervation regulates epithelial secretion of CSF (Lindvall et al., 1978; Shipley *et al*., 2020) as delivery of serotonin agonists triggers coordinated waves of calcium activity on epithelial cells that culminated in apocrine CSF secretion (Shipley *et al*., 2020). Nevertheless, the question remains on whether epithelial cells specialize in receiving and propagating neuronal signals (such as through tight junctions) or, alternatively, exhibit varying response threshold to neurotransmission. We have identified a subtype of epithelial cells within the lateral ventricle CP, which may carry out specific functions. Our findings suggest that *Atp10b*^+^/*Gata3*^+^ epithelial subtype might be sensitive to neurotransmission and could be the first responders to neuronal input-triggered apocrine secretion.

The presence of ‘ciliogenic’ epithelial cells was previously reported at embryonic stages (Dani *et al*., 2021). Here we observed this population at neonatal time points indicating an ongoing process of cell maturation at postnatal stages. We further demonstrate a distinct complexity of lateral ventricle CP cells over time by flow cytometry. A very low proportion of proliferating epithelial cells (not ‘ciliogenic’) was detected both in neonatal and adult stages. These results are in line with previous studies (Dani *et al*., 2021; Liddelow et al., 2010). Importantly, genes involved in ciliogenesis and cell division exhibit chromatin accessibility across all epithelial subtypes, suggesting that mature epithelial cells may re-engage in these processes, supporting their renewal, as observed in epithelial tissues of other organs.

Despite the lack of confirmation of a *Folr1/Cp/Gpx3* triple positive cells in the tissue, epithelial cells expressing high levels *Gpx3* were easily distinguishable in lateral ventricle CP. The expression of *Cp* and *Gpx3* in the CP was described before (Aldred et al., 1987; Klomp et al., 1996; Kratzer et al., 2013; Planques et al., 2021; Yang et al., 2021), but not associated. Furthermore, a Gpx3^high^ epithelial cell population was recently described in human lateral ventricle CP by single nucleus RNA-seq (Yang *et al*., 2021), and induction of Gpx3 was observed in response to LPS (Marques et al., 2009b).

Our study further reveals the presence of open chromatin in immune-response genes within epithelial cells, despite their lack of expression, suggesting that a subset of epithelial cells exhibit chromatin priming for inflammation-responsive genes. This allows lateral ventricle CP epithelial cells to mount a rapid response, occurring as early as 1 hour post-challenge (Marques *et al*., 2009a). Interestingly, these genes are also described to play a role in multiple sclerosis (MS). CCL2 and CXCL1 have been reported to be dysregulated in MS (Ghafouri-Fard et al., 2021), STEAP4 was reported to be a key Th17-specific effector molecule required for the initiation and pathogenesis of the Experimental Autoimmune Encephalomyelitis (EAE) MS mouse model (Zhao et al., 2021) and blocking CSF-1/CSF-1R ameliorated MS pathology in EAE (Hwang et al., 2022). Whether these genes are dysregulated in the lateral ventricle CP of MS remains an open question.

A chronic low-grade activation of the innate immune system, inflammaging, has been documented to impact the brain parenchyma immune cells, microglia (Costa et al., 2021). An age-related immune phenotype was also observed in the CP (Baruch *et al*., 2014; Dani *et al*., 2021; Mesquita *et al*., 2015). Remarkably, we found inflammation markers and host-defense ligand-receptor interactions in CP cell types starting at neonatal stages and gradually increasing to young adult stages. Strikingly, the increase in immune response genes was not confined to macrophages but comprised epithelial, fibroblasts and endothelial cells indicating a widespread inflammaging. Immune cells of the lateral ventricle CP were more abundant in adult and exhibited a different cell-type profile when compared to postnatal stages. In the process of validating markers for epiplexus cells, we identified an epiplexus cell bridging the lateral ventricle CP and brain parenchyma, suggesting that these cells might actively travel between the brain and the lateral ventricle CP, with implications for health and disease that remain unexplored.

In summary, by mapping the transcriptome and chromatin accessibility of lateral ventricle CP cells over time, we offer detailed cell profiles that will be essential for assessing their potential in targeted therapies, particularly by harnessing the plasticity of epithelial cells.

## METHODS

### Animals Mice

For single-cell multiome, immunofluorescence and RNAscope in situ hybridization we used C57BL/6J mice purchased from Charles River laboratories, C57BL/6NTac-Ttr^tm1(EGFP/cre/ERT2)Wtsi^ (EMMA ID EM:09763) purchased from the INFRAFRONTIER/EMMA partner Mary Lyon Centre at MRC Harwell.

Sox10:Cre-RCE:LoxP strains used for RNAscope in situ hybridization, were purchased from The Jackson Laboratory: *Gt(ROSA)26Sor^tm1.1(CAG-EGFP)Fsh^*/Mmjax (RRID:MMRRC_032037-JAX) and B6;CBA-Tg(Sox10-cre)1Wdr/J (IMSR_JAX:025807) .

Adult mice were housed in groups of 2–6, breeding cages were comprised of 1 male and up to 2 females. Mice were maintained under the following standard laboratory conditions: 12-hour light/dark cycle (lights on from 8:00 to 20:00), temperature 22 °C ± 1 °C and relative humidity 60%. A standard diet and water ad libitum was provided. To ensure specified pathogen-free health status, regular testes for pathogens were performed in sentinel mouse housed in the same room. Animal procedures and experimenters were certified by the Portuguese regulatory entity, Direção Geral da Alimentação e Veterinária (DGAV), approved by the Ethics Committee of the Life and Health Sciences Research Institute and by DGAV (003081), and conducted in accordance with European Union regulations (directive 2010/63/EU) on animal care and experimentation.

For single-cell RNA-seq experiments adult and pregnant C57BL/6J mice were purchased from Charles River laboratories. Adult mice were house at a maximum of 5 adult mice per IVC-cage of type II Allentown and pregnant mice were house separately. Mice were maintained under a pathogen-free environment at the animal facility under the following standard laboratory conditions: light/dark cycle (dawn 6:00-7:00; daylight 07:00-18:00; dusk 18:00-19:00; night 19:00-06:00), temperature 22 °C and consistent relative air humidity of 50%. All experimental procedures were performed following the guidelines and recommendations of local animal protection legislation and were approved by the local committee for ethical experiments on laboratory animals (Stockholms Norra Djurförsöksetiska nämnd in Sweden), lic nr. 1995/2019.

### Rats

Wistar Han rats (Rattus norvegicus) were purchased from Charles River (France). Adult rats were housed in groups of 2–3, breeding cages were comprised of 1 male and up to 2 females. Rats were maintained under the following standard laboratory conditions: 12-hour light/dark cycle (lights on from 8:00 to 20:00), temperature 22 °C ± 1 °C and relative humidity 60%. A standard diet and water ad libitum was provided. To ensure specified pathogen-free health status, regular tests for pathogens were performed in sentinel mice housed in the same room. Animal procedures and experimenters were approved by the Ethics Committee of the Life and Health Sciences Research Institute and by DGAV (DGV9457) and conducted in accordance with European Union regulations (directive 2010/63/EU) on animal care and experimentation.

### Genotyping Ttr^tm1(EGFP/cre/ERT2)Wtsi^ mice

Ear biopsies were collected during weaning between 21 to 26 days of age. DNA extraction was performed by incubating ear biopsies in 75ul of NaOH 25mM/EDTA 0.2mM for 1h at 98°C followed by neutralization with 75ul of Tris HCl 40mM pH5.5. After centrifugation for 3min at 4000 rpm, 1µl of supernatant was used for PCR using DreamTaq Green PCR Master Mix 2x (K1081, ThermoFisher) according to the manufactorer’s instructions. Primers sequences were provided by the provider and consisted of 3 primers: CAS_R1_Term TCGTGGTATCGTTATGCGCC, Ttr_377681_F CCTTAGGCCATGGGACACT and Ttr_377681_R2 GATTCAATCCCCAACAACG. PCR comprising primers Ttr_377681_F and Ttr_377681_R2 resulted in a wild type band of 454bp, Ttr_377681_F and CAS_R1_Term resulted in a mutant band of 89bp, that were further ran in a 2% agarose gel and analysed under a ChemiDoc (Biorad).

### Tissue collection and processing for RNA-seq

Bulk RNA-seq from adult male rats comprised 5 main timepoints: postnatal day (P) P1, P4, P7, P10 and P60. Animals were anesthetized with pentobarbital (200mg/kg) and were transcardially perfused with sterile RNase free cold saline (0.9% NaCl). Brains were removed and the choroid plexuses (CP) from the lateral ventricles were isolated under a stereo microscope (SZX7; Olympus) with fine forceps, frozen in dry ice, and stored at -80°C.

### RNA extraction and quality analysis for RNA-seq

Total RNA extraction was performed using SPLIT RNA Extraction Kit (Lexogen, Viena, Austria). Briefly, the CPs were homogenized in an isolation buffer highly chaotropic. Samples, were then transferred to a phase lock gel column, which contains a gel matrix that, based on the density differences, acts as a barrier between the organic and aqueous phase. Acidic phenol and acidic buffer were added to create a monophasic solution, which allows the separation of genomic DNA into the organic phase. Chloroform was added and the phases were separated by centrifugation for 2min at 12.000xg at 18°C. The RNA, contained in the aqueous phase, was decanted into a new micro tube. The RNA was then precipitated onto a silica column by addition of 1.75x volume of isopropanol. The RNA was further purified by washing the column. Finally, RNA was eluted in 20μL of Elution Buffer. Quality of total RNA extracted was evaluated using Experion RNA StdSenses Analysis Kit (Bio-Rad, Hercules, California, USA). Briefly, heat denaturated RNA was loaded into an electrophoresis system and detected measuring fluorescence of a fluorophore that binds to RNA. RNA quality is measured by the RNA quality index (RQI). Using an algorithm, RQI measures RNA integrity comparing the electropherogram of RNA samples to a series of standardized degraded RNA samples. The RQI method returns a number between 10 (intact RNA) and 1 (highly degraded RNA).

### Tissue dissociation for single-cell RNA-seq, single-cell multiome and flow cytometry

Single-cell RNA-seq comprised three main timepoints: postnatal day (P) P2-3, P11 and 10 week old young adult mice. Single-cell multiome samples comprised two time points: P11 and young adult mice (10-14 weeks). Animals were anesthetized and transcardially perfused with cold PBS without Ca^2+^ and Mg^2+^ (10010023, ThermoFisher). Brains were removed and the choroid plexuses (CP) from the lateral ventricles were isolated under a stereo microscope with fine forceps and dissected into small pieces with a high precision dissecting micro scissor. A pool of 3-4 CP from the lateral ventricles of both female and male mice were dissociated to acquire one sample. CPs were further dissociated in 0.1 mg/ml pronase (10165921001, Sigma) and 20U/ml of DNase I (4536282001, Sigma) in 2mL of PBS for 15 min in the water bath at 37°C, shaken every 5min or less. After centrifuging at 400g for 5min, supernatant was discarded and pellets were resuspended in 50µl of enzyme P + 950µl of buffer Z and 10µl buffer Y + 5µl of enzyme (adult brain dissociation kit, 130-107-677, Miltenyi Biotec) followed by an incubation for 30 min at 37°C and a mechanic digestion consisting of pipetting up and down 20x with P1000 every 5 min. Samples were then centrifuged for 5min at 400g, supernatant was discarded and pellets were resuspended in 1ml of trypsin 0.025% diluted from a stock solution of 0.25% trypsin-EDTA (25200056, ThermoFisher) in PBS without Ca^2+^ and Mg^2+^. Upon an incubation of 15min at 37°C, with mechanic digestion as described above, the samples were passed through 30µm strainer (CellTrics® strainers, Sysmex partec) pre-wet with 1ml of PBS-2%BSA (A8412, Sigma) and washed in 7 ml of PBS-2%BSA. After centrifugation at 400g for 5min pellets were resuspended in 1ml of PBS-2%BSA and passed through a 20µm strainer (CellTrics® strainers, Sysmex partec), pre-wet with 0.5ml of PBS-2%BSA washed in 1ml of PBS-2%BSA. Lastly, samples were centrifuged at 400g for 5min and resuspended in 40µl of PBS-2%BSA for further cell counting with trypan blue. The Chromium Next GEM Single Cell 3ʹ Reagent Kits v3.1 from 10x genomics were used for single-cell RNA-seq, and protocols were followed according to the user guide.

For the single-cell multiome, cells were extracted as described above and comprised two timepoints: postnatal day P11 and approximately P60 young adult mice. Nuclei isolation was as follows: cells were resuspended in chilled lysis buffer (containing 0.01% IGEPAL (CA-630), 0.01% Tween-20, 0.001% Digitonin, 1% BSA, DTT 200mM, 10 mM Tris-HCl pH 7.4, 10 mM NaCl, 3 mM MgCl2 and RNAse inhibitor (AM2684, Ambion ThermoFisher)) and incubated on ice for 3 min. After the incubation, wash buffer (containing 0.1% Tween-20, 1% BSA, DTT 200mM, 10 mM Tris-HCl pH 7.4, 10 mM NaCl, 3 mM MgCl2 and RNAse inhibitor 1U/µl) was added on top without mixing and the nuclei were centrifuged for 5 min at 500 g and 4°C. An extra isolation step was performed by adding 100 µl of with Nuclei EZ lysis buffer (Sigma, #N3408) to the pellets and pipetting up and down 5x followed by centrifugation at 500g for 5min at 4°C. Supernatant was discarded and pellets were resuspended in chilled lysis buffer followed by a washing step as described above. Lastly, nuclei were washed once in Diluted Nuclei buffer (10x Genomics) containing 1% BSA, DTT 20mM and RNase inhibitors 1U/µl, centrifuges at 500g for 5min at 4°C and resuspended in 10µl of diluted nuclei buffer (10x genomics). 2µl of nuclei were counted with trypan blue and the rest was used for tagmentation according to 10x genomics multiome protocol. Chromium Next GEM Single Cell Multiome ATAC + Gene Expression kits from 10x genomics were used for the single-cell multiome. Protocols were followed according to the user guide.

### Flow cytometry analysis

For flow cytometry, cells were extracted as described above and comprised two timepoints: postnatal day P11 and P60 young adult mice. At the final wash, cells were resuspended in 50 μL of anti-CD45 Fitc (clone HI30, Biolegend; 1:100 dilution in FACS buffer - PBS, 2% BSA, 0.01% Sodium azide and 2mM Na2EDTA) and incubated for 20 min in the dark at room temperature. Afterwards, cells were washed, resuspended in FACS buffer with 7-AAD (0.167 μg/mL) and incubated for 10 min before being acquired in a LSRII flow cytometer using the FACS Diva Software v6.0 (Becton Dickinson, NJ, USA). Data were analyzed in a blinded way, using the FlowJo Software v10 (Becton Dickinson, NJ, USA).

### Tissue collection and processing for immunofluorescence and RNAscope in situ hybridization (ISH)

Mice were anesthetized and transcardially perfused with cold PBS and 4% paraformaldehyde in PBS. Brains were collected and transferred to tubes with 4% paraformaldehyde. After 1h of post fixation the solution was replaced by 30% sucrose in PBS and kept at 4°C until the brains sink. Brains were then embedded in tissue-teck O.C.T. compound, frozen in dry ice, and kept at -80°C for further cryosectioning. Coronal slices of 20µm comprising the lateral ventricle CP were collected on a cryostat (Leica CM 1950) and frozen at -80°C.

For free floating RNAsope in situ hybridization and immunofluorescence, CP was collected from fixed brains under a stereo microscope and further post-fixed for 30 mins and used immediately for subsequent procedures.

### RNAscope ISH and immunofluorescence

RNAscope ISH was performed as described previously (Falcão et al., 2018) on brain sections and free-floating CPs with the following probes for mouse *Folr1*, Otx2, *Gata3*, *Atp10b*, *Ptprc, Sox10, Izmb*, *Cdk1*, *Pecam1, Ccno, Syngr, Myl9* and *Lum* all purchased from ACD. RNAscope ISH protocol for sections was performed following manufacturer’s instructions with minor modifications (ACD, RNAscope® Multiplex Fluorescent Detection Reagents v2, 323110). Briefly, sections and free-floating CPs were placed on a hot plate (100 °C) with 1x target retrieval reagent (322000) for 5 min followed by 2 steps of washes of 2 min and 1 wash with 100% ethanol for 2 min. Protease IV was applied on top of the sections and incubated for 20 min at room temperature, followed by 2 washes of 2 min each. Probes were diluted 1:50 in the C1 probe, hybridized for 2 h at 40 °C and washed twice in wash buffer (310091). Amplification steps were performed by incubating with v2Amp1 (30 min), v2Amp2 (30 min) and v2Amp3 (15 min) at 40 °C with washes of 2 × 2 min in between steps. Sections were incubated with v2-HRP-C1 for 15 min at 40 °C and washed twice in wash buffer for 2 min. Opal fluorophores from Akoya Biosciences (Opal 520, PN FP1487001KT; Opal 570, PN FP1488001KT; Opal 690, PN FP1497001KT) were diluted 1:1500 in TSA buffer (322809) and incubated for 30 min at 40 °C followed by 2 washes of 2 min and HRP blocker incubation for 30 min at 40 °C. The last steps were performed subsequently for v2-HRP-C2 and v2-HRP-C3. Lastly, sections and free-floating CP were incubated 2-10 min in DAPI (D1306, ThermoFisher), washed twice and mounted in either Fluoromount-G (00-4958-02, ThermoFisher) or ProLong Gold Antifade Mountant (Invitrogen Cat# P36934) and kept at 4 °C until further microscopic analysis.

For immunofluorescence the following antibodies were used: MOG (MAB5680, Sigma, 1:100), ß3-tubulin (g712a, Promega 1:100), CNP (ab6319, Abcam, 1:200), NFH (ab8135, Abcam 1:1000).

Free floating CPs were incubated overnight at 4°C with primary antibodies diluted in PBS/0.5% Triton/10% normal donkey serum (D9663, Sigma). After washing with PBS, secondary Alexa Fluor-conjugated antibodies (Invitrogen) diluted 1:1000 in PBS/0.5% Triton/10% normal donkey serum were incubated for 2 h at room temperature. CPs were washed twice in PBS and incubated 2 min in DAPI. Lastly, CPs were mounted in Superfrost Plus™ slides in Fluoromount and kept at 4 °C until further microscopic analysis.

### Confocal microscopy

Images were collected by confocal microscopy (FLUOVIEW FV1000 and FV3000, Olympus) and analyzed in Image J.

### Bulk RNA-seq QC and differential gene expression

The bulk RNA-seq samples were preprocessed for adapter/quality trimming and then aligned to the rattus norvegicus transcriptome using STAR (Dobin et al., 2013) version 2.7 – quantMode –sjdbOverhang 99 with EnsEMBLv75 gtf annotations. Only uniquely mapped reads were retained for downstream analysis using SAMtools (Danecek et al., 2021). Aligned samples were converted to bedgraph files using Deeptools (Ramírez et al., 2016) bamcoverage for each strand and normalized to total of reads. Filtered fastq files were used in Salmon 0.8.2 (Patro et al., 2017) to recover the raw reads counts and transcript per million (TPM) values per transcript and gene. The differential gene expression analysis was performed with Deseq2 (Love et al., 2014). Results from differential expression were plot using Enhancedvolcano package with log2 fold change and adjusted p value from Deseq2.

### scRNA-seq 10X Genomics preprocessing

Our dataset is composed of eight samples: one lateral ventricle CP at Postnatal day 2 (P2), two at P3, three at P11, and three at P60. scRNA-seq data from 10X Genomics was processed with the 10x Genomics Cell Ranger v5.0.1 *count* function using default parameters. The reads were aligned to mm10 reference genome for each sample. Then, using Scanpy package (Wolf et al., 2018) (version 1.9.8) all the samples were combined, and the data was log normalized. Doublet detection and removal were then performed using scrublet (Wolock et al., 2019) (version 0.2.3) with the threshold set to 0.33. Data filtering was then performed by removing all the cells that had less than 250 genes, all the genes that were expressed in less than 5 cells and using the following metrics: total_counts < 80000, n_genes_by_counts < 9000, pct_counts_Mito < 20, ct_counts_Hb < 0.05, pct_counts_Ribo < 20, pct_counts_Ribo > 2.5. This resulted in a total of 47550 cells.

### scRNA-seq Dimensionality reduction and clustering

Dimensionality reduction and clustering were performed using Scanpy (version 1.9.8). First highly variable gene (HVG) selection was performed, keeping the first 4000 more variable genes using the function *highly_variable_genes* with the parameter *flavor=’seurat_v3’* and *n_top_genes=4000.* Then PCA was performed and the first 32 PCs were kept to perform clustering and data visualization. UMAP was calculated using the functions *sc.pp.neighbors* with the *n_pcs=32* and *n_neighbors=15* and *sc.tl.umap* with default parameters. Data integration was also performed using Harmony (Korsunsky et al., 2019) (version 0.1.8) to reduce the batch effect between the different samples. Then clustering was performed using the Leiden algorithm accessed by the function *sc.tl.leiden.* Marker genes were identified by using the function *sc.tl.rank_genes_groups* using the parameters *method=’wilcoxon’* and *use_raw=True*. From this analysis resulted 5 clusters corresponding to the 5 major cell types found (Epithelial cells, Endothelial cells, Mesenchymal cells, Immune cells, Neuronal/Glial cells). While performing subclustering in the different cell types, we were able to remove more doublets using scrublet and manually removing clusters that presented several markers for at least two different cell types, a high number of counts per cell, and high number of genes. After subclustering, the main dataset was re-filtered to include only the cells retained in the final subclusters of each cell type, resulting in a total of 41,372 cells.

### Cell type proportions analysis

After cluster and cell type annotation we calculated cell proportions for each cell type considering the three different time points. For that we used the function pd.crosstab from the python package Pandas (McKinney, 2010) as follows pd.crosstab(adata.obs[’batch’], adata.obs[’Cell_type’], normalize=’index’). This allowed us to create a frequency table for cell type proportions normalized within each cell type.

### scRNA-seq Differential expression analysis

Differential expression (DE) was performed using Scanpy (version 1.9.8). First, we divided the filtered raw data into 4 subsets: macrophages, fibroblasts, endothelial cells, and epithelial cells. Proliferating cells from these groups were not considered in this analysis and each subset was treated independently. Then we randomly selected an equal number of cells from each age group, 1100 cells/group for epithelial cells, 320 cells/group for endothelial cells, 680 cells/group for fibroblast, and 180 cells/group for macrophages. Then the data was log normalized and scaled using the functions *sc.pp.normalize_per_cell*, *sc.pp.log1p*, *sc.pp.scale*, respectively, with default parameters except the first one which had the parameter *counts_per_cell_after=1e4*. Then we used the function *sc.tl.rank_genes_groups* with parameters *method=’wilcoxon’* and *use_raw=False*, to identify the DE genes of each age group. The parameters *groups* and *reference* were used to select which age groups were being used to calculate the DE genes and which one was the reference. Here all age groups were tested against each other in groups of two using the different combinations. Selected genes had an adjusted p-value < 0.05 and a log foldchange > 0.5. The cutoff used for gene ontology analysis was a log foldchange of 1.9.

### Cell-Cell Communication

Cell-Cell communication inference was performed using scRNA-seq data at two different time points, P2/P3 and P60. It was calculated within each time point using Liana package (Dimitrov *et al*., 2022) (version 0.1.14) and the function *liana_wrap* with the parameter *resource = ’MouseConsensus’.* Then, using *liana_aggregate* results were combined and summarized. To get unique ligand-receptor pairs for each pair of cell types we sorted our results by NATMI’s edge specificity (Hou et al., 2020), selected the first 40 ligand-receptor pairs, and plotted the results using *liana_dotplot.* To get the interaction frequencies for the pair of cell types that could potentially communicate, we filtered our results to include only the interactions that were found between all methods on the consensus, resulting interactions were plotted using *heat_freq*.

### Single-cell multiome 10X Genomics preprocessing

Single-cell multiome data from 10X Genomics was processed with the 10x Genomics Cell Ranger ARC v1.0.0 *count* function using default parameters. The reads were aligned to a custom mm10 reference genome, to include the eGFP and CreERT2 genes. Our data is composed of 6 samples, 3 P11, and 3 P60. Each sample was processed with CellRanger ARC individually. One of the P60 had the expression of the Ttr gene replaced by the expression of eGFP. Then, using Seurat package (Hao et al., 2024) (version 5.1.0) all the samples were combined, and the data was log normalized. For snRNA-seq data, doublets detection and removal were then performed using DoubletFinder (McGinnis et al., 2019) (version 2.0.4) separately for each sample . For snATAC-seq data we used AMULET (Thibodeau et al., 2021) (version 1.1) with default parameters. Data filtering was performed individually for each sample (QC presented in Fig S2), nucleossome_signal ≤ 2.0, and TSS.enrichment ≥ 2.0 were applied for all samples. This pre-processing resulted in a total of 39079 cells. Afterwards we performed peak calling using MACS2 (Gaspar, 2018) within Signac (Stuart et al., 2021) (version 1.14.0) to retrieve a consistent set of peaks over the entire dataset.

### Single-cell multiome Dimensionality reduction, clustering and cell type annotation

The following analysis was performed using both Seurat (version 5.0.1) and Signac (version 1.14.0) for the corresponding modality. For snRNA-seq data, HVG selection was performed, keeping the first 4000 more variable genes using the function *FindVariableFeatures* with parameters *selection.method = “vst”, nfeatures=4000.* Then PCA was performed and the first 20 PCs were kept followed by clustering and data visualization. For snATAC-seq data we performed Latent Semantic Indexing (LSI) with dims = 2:30. Following that, Harmony (version 1.2.3) was used in both modalities based on the sample variable and using 1:50 PCs for RNA modality and 2:50 for ATAC modality. UMAP was calculated using the function *RunUMAP* with *dims = 1:20* and *n.neighbors = 15* for RNA modality and *dims = 2:30* for ATAC modality. UMAP visualization was performed using *DimPlot* for both cases. Clustering was performed using *FindNeighbors* with *dims=1:20* for RNA and *dims=2:30* for ATAC modality., followed by *FindClsuters* in both modalities. Marker genes were identified by using the function *FindAllMarkers* using the parameters *test.use=“wilcox”* and *only.pos=TRUE*.

Marker peaks were identified by using the function *FindAllMarkers* using the parameters *test.use = ’wilcox’* and *min.pct = 0.1.* Cell type annotation resulted in 5 different cell types (Endothelial cells, Epithelial cells, Immune cells, Mesenchymal cells, and Neuronal and Glial cells). We also used *transfer_*anchors using our scRNA-seq data to double check cells identities found here. Subclustering was performed on each cell type, using the same methods described above. QC metrics used in Figs. S3 to S7. Joint graph visualization of both modalities was achieved using the weighted nearest neighbor (WNN) from FindMultiModalNeighbors with *dims.list=list(1:20, 2:30)*. While performing subclustering in the different cell types, we were able to remove more doublets using scrublet and manually removing clusters that presented several markers for at least two different cell types, a high number of counts per cell, and high number of genes. After subclustering, the main dataset was re-filtered to include only the cells retained in the final subclusters of each cell type, resulting in a total of 53835 cells.

### Peak annotation

Peaks were annotated using HOMER (Heinz et al., 2010) (version 5.1) function annotatepeaks.pl. This annotation was then used to perform the donut plots with the peaks annotation of the different cell types.

### Gene Ontology analysis

GO analysis was performed in ShinyGO 0.80: a graphical gene-set enrichment tool for animals and plants (Ge et al., 2020) with the following settings: GO Biological process and Cellular component, FDR cutoff: 0.05, minimum number of genes: 3. Pathways selected with lowest FDR values and redundancy removal.

## Data availability

Raw data is deposited in GEO with the following accession numbers: RNAbulk-seq data GSE296843, scRNA-seq data GSE296842, and scMultiome-seq data GSE296841.

The scRNA-seq and scMultiome dataset will soon be available and can be explored at the web resource Cell Browser (https://cells.ucsc.edu) (Speir et al., 2021).

## AUTHOR CONTRIBUTIONS

A.M.F., J.C.S., M.F. and L.L. conceived the idea and designed the study. A.M.F, E.C. and M.M performed single-cell RNA-seq and multiome sample preparation. F.M, D.A. and J.C.S. performed RNA-seq sample preparation. M.F. and L.L. performed all computational data analysis with the help of A.M.F, E.A. and G.C.B. A.M.F., A.V., J.C, A.I.A and N.M. have performed validation experiments. A.M.F. and C.N. performed the flow cytometry experiments. A.M.F. wrote the manuscript with the help of all authors. All authors read and approved the final version of the manuscript.

## ACKNOWLEDGMENTS

The authors acknowledge support from the National Genomics Infrastructure in Stockholm funded by Science for Life Laboratory, the Knut and Alice Wallenberg Foundation and the Swedish Research Council. Computation/data handling was enabled by resources provided by the National Academic Infrastructure for Supercomputing in Sweden (NAISS), partially funded by the Swedish Research Council through grant agreement no. 2022-06725, and by Portuguese Foundation for Science and Technology (https://doi.org/10.54499/2024.08336.CPCA.A0).

The authors acknowledge the Eukaryotic Single Cell Genomics (ESCG) facility in Stockholm funded by Science for Life Laboratory, KI Core and StratRegen. This work was also supported by the ICVS Scientific Microscopy Platform, a member of the national infrastructure PPBI— Portuguese Platform of Bioimaging (PPBI-POCI-01–0145-FEDER-022122).

We thank the Wellcome Trust Sanger Institute Mouse Genetics Project (Sanger MGP) and its funders for providing the mutant mouse line (Allele: Ttr^tm1(EGFP/cre/ERT2)Wtsi^), and INFRAFRONTIER/EMMA (www.infrafrontier.eu) partner [Mary Lyon Centre at MRC Harwell] from which the mouse line was received. Funding information may be found at www.sanger.ac.uk/mouseportal and associated primary phenotypic information at www.mousephenotype.org. Mouse services were provided by the Mary Lyon Centre at MRC Harwell (www.har.mrc.ac.uk).

Funding: A.M.F. was supported by Portuguese Foundation for Science and Technology (grant no. 2020.02753.CEECIND) and by a fellowship from “la Caixa” Foundation (ID100010434) with the fellowship code LCF/BQ/PI19/11690005. This work was funded by Swedish Research Council (grant no. 2019-02030), Bial Foundation Grant (217/12) and by national funds, through the Foundation for Science and Technology (FCT), under projects UID/06304/2023 and LA/P/0050/2020 (DOI 10.54499/LA/P/0050/2020).

## Non-author contributions

We would like to thank Ruben Silva, Diogo Monteiro and Miguel Pacheco, at the time master students that participated and provided technical assistance. We thank Prof Nuno Sousa for supporting and helping in the establishment of single-cell RNA-seq technologies at ICVS, Prof Margarida Correia-Neves for the helpful advice and discussions of the data, Prof Joana Palha for advice and proof-reading, and Prof António Salgado for providing antibodies. We are grateful to Tony Gimenez-Beristain, Chao Zheng for providing assistance with animal work and laboratory support.

