## Supplementary Figures for "Multi-omics Profiling of the Lateral Ventricle Choroid Plexus Reveals Developmental Cellular Remodeling, Early Immune Gene Activation, and a Novel Epithelial Subtype"

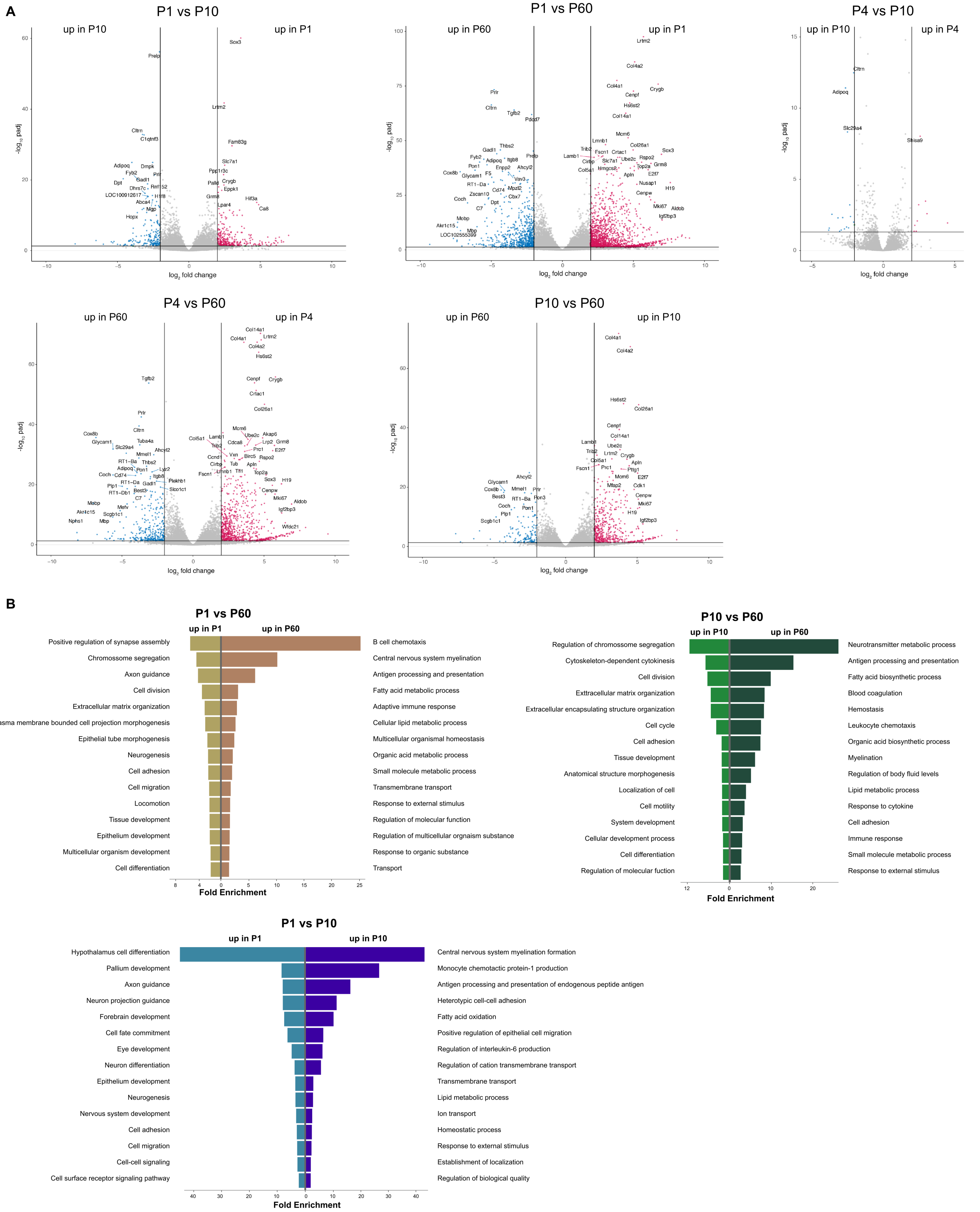


**Figure S1:Bulk RNA-seq of rat LVCP at different postnatal stages and adult.**

(A) Volcano plots representing differencial gene expression in RNA-seq from rat LVCP from different time points. n=3 for P1, P4 and P10, n=2 for P60. Genes with adj. p value < 0.05 and log2 fold change >2 are shown in blue for later time points and red for earlier time points.

(B) Gene ontology (GO) analysis for the biological function og genes differentially expressed in the different time points.

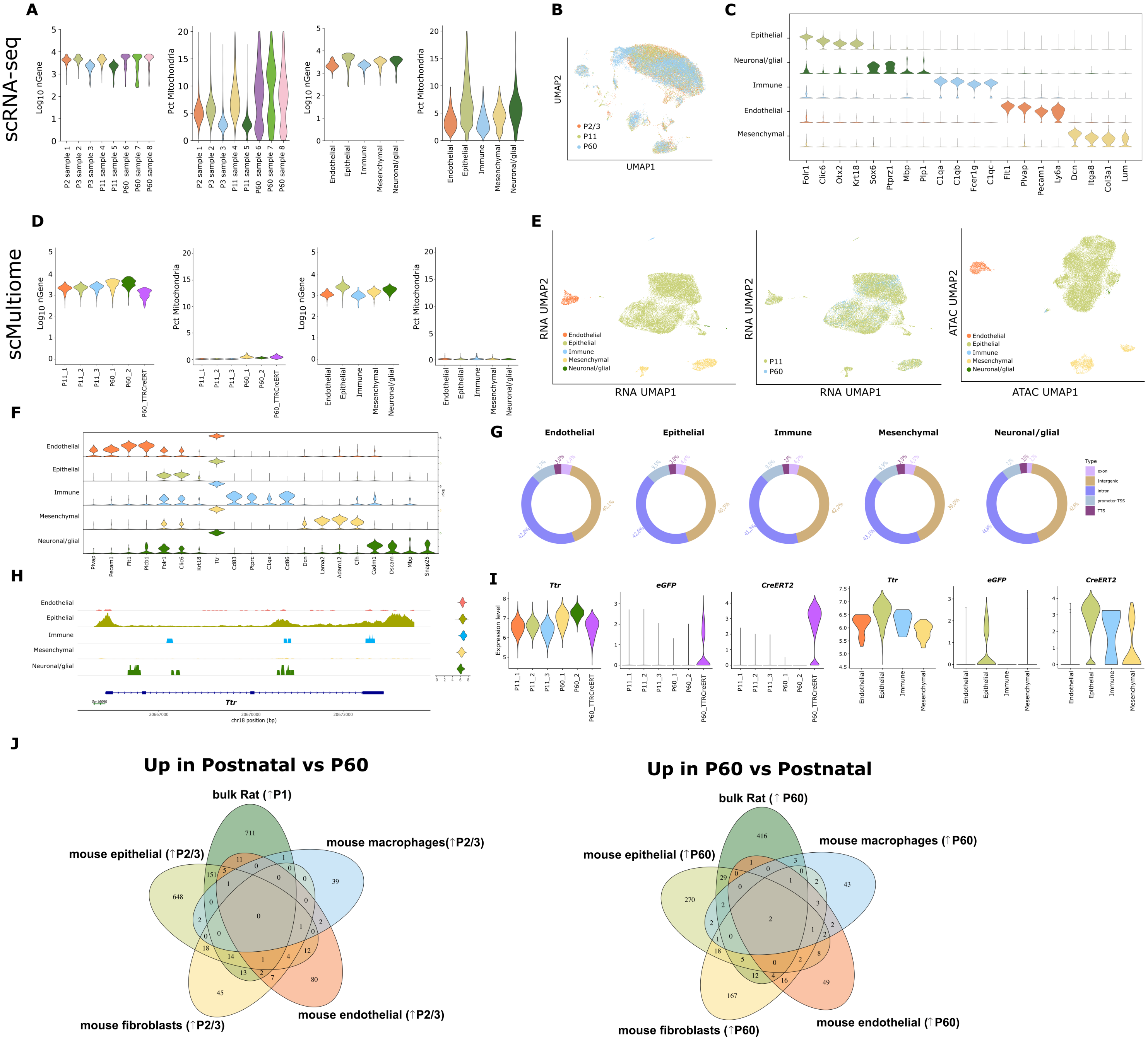


**Figure S2: scRNA-seq and scMultiome QC analysis and cell markers, overlapp of differencially expressed genes between postnatal and adults in different species.**

(A) Violin plots representing QC analysis for number of genes (log10nGene) and percentage (pct) of mithocondrial genes, color-coded by sample and by cell-type in the scRNA-seq data.

(B) UMAP plot of all cells analysed by scRNA-seq, color-coded by time point.

(C) Violin plot of median gene expression of four markers per cell population in the scRNA-seq data.

(D) Violin plots representing QC analysis for number of genes (log10nGene) and percentage (pct) of mithocondrial genes, color-coded by sample and by cell-type in the scMultiome data.

(E) UMAP plot of all cells analysed by scMultiome, for RNA and ATAC, color-coded by sample type and time point.

(F) Violin plot of median gene expression of four markers per cell population in the scMultiome data.

(G) Distribution of the differentially accessible peaks amongst cell popupations.

(H) IGV tracks of CA showing *Ttr* chromatin accessibility amongst cell-types identified in scMultiome. Violin plots depicting the expression of the genes, and the corresponding genomic coordinates are shown..

(I) Violin plots of expression level of *Ttr*, *eGFP* and *CreERT2* amonsgt samples and cell types.

(J) Venn diagram representating the overlapp of genes up-regulated in postnatal stages versus adult P60 of both rat bulk RNA-seq and mouse scRNA-seq. Compari-sons perfomed: up-regulated in P1vs P60 of rat bulk RNA-seq, and up-regulated in P60 vs P2/3 of mouse scRNA-seq descriminated by the four main lateral ventricle CP cell-types.

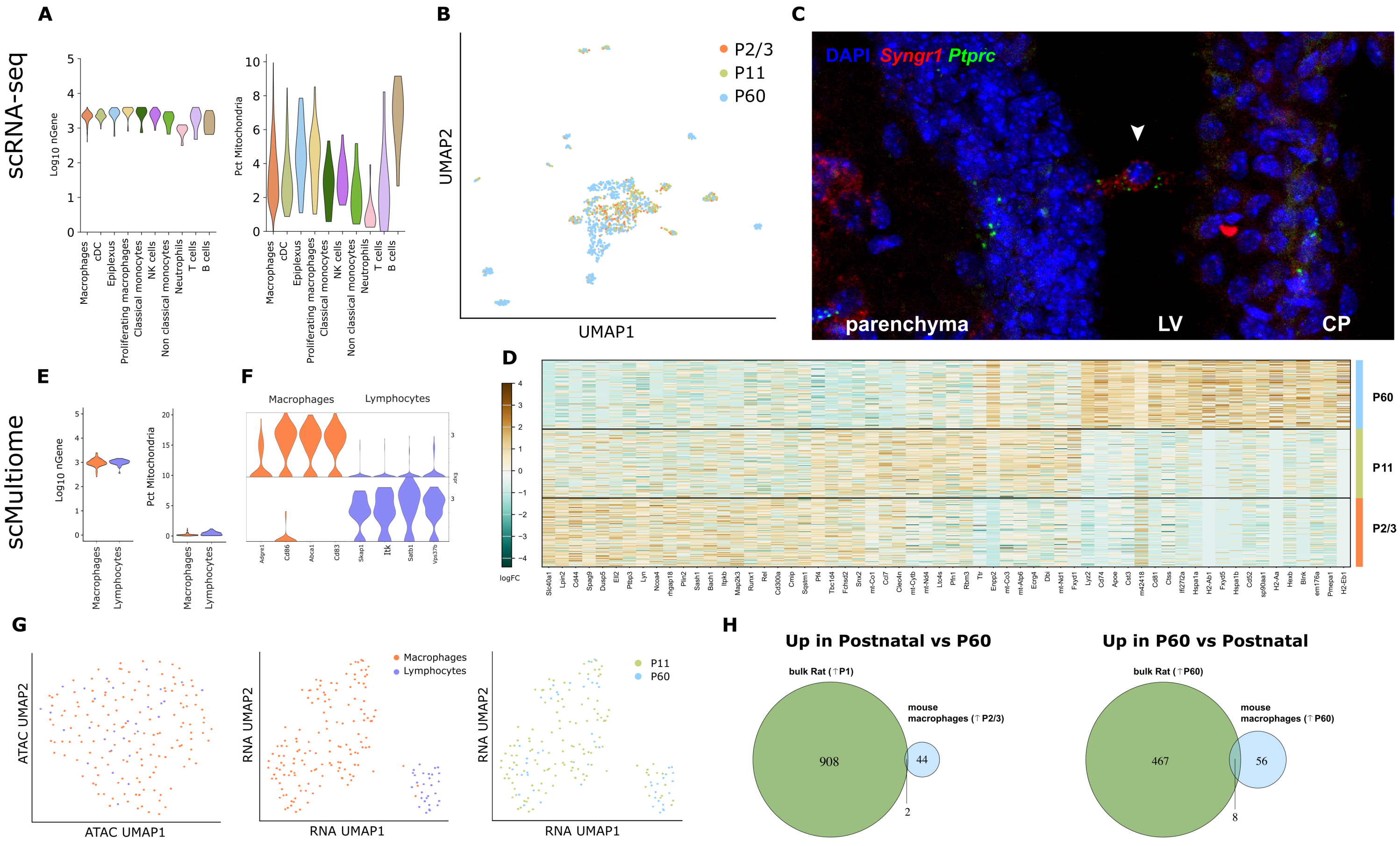


**Figure S3: scRNA-seq and scMultiome QC analysis for immune cells, and overlapp of differencially expressed genes between postnatal and adults in different species**

(A) Violin plots representing QC analysis of the 10 immune cell populations in scRNA-seq, number of genes (log10nGene) and percentage (pct) of mithocondrial genes color-coded by cell-type.

(B) UMAP plot of immune cells analysed by scRNA-seq, color-coded by time point.

(C) RNAscope ISH representing a tissue section of lateral ventricle CP marked with probes for *Ptprc* (immune cell marker) and *Syngr1* (epiplexus cell marker). P60 mouse lateral ventricle CP.

(D) Heatmap of differential expression genes in macrophages across different ages.

(E) Violin plots representing QC analysis of the two immune cell populations in scMultiome, number of genes (log10nGene) and percentage (pct) of mithocondrial genes color-coded by cell-type.

(F) Violin plot of median gene expression of four markers per cell population in the scMultiome data.

(G) UMAP plots of immune cells analysed by scMulti-ome, for RNA and ATAC, color-coded by sample type and time point.

(H) Venn diagram representating the overlapp of genes up-regulated in postnatal stages versus adult P60 (and vice-versa) of both rat bulk RNA-seq and mouse immune cells from scRNA-seq.

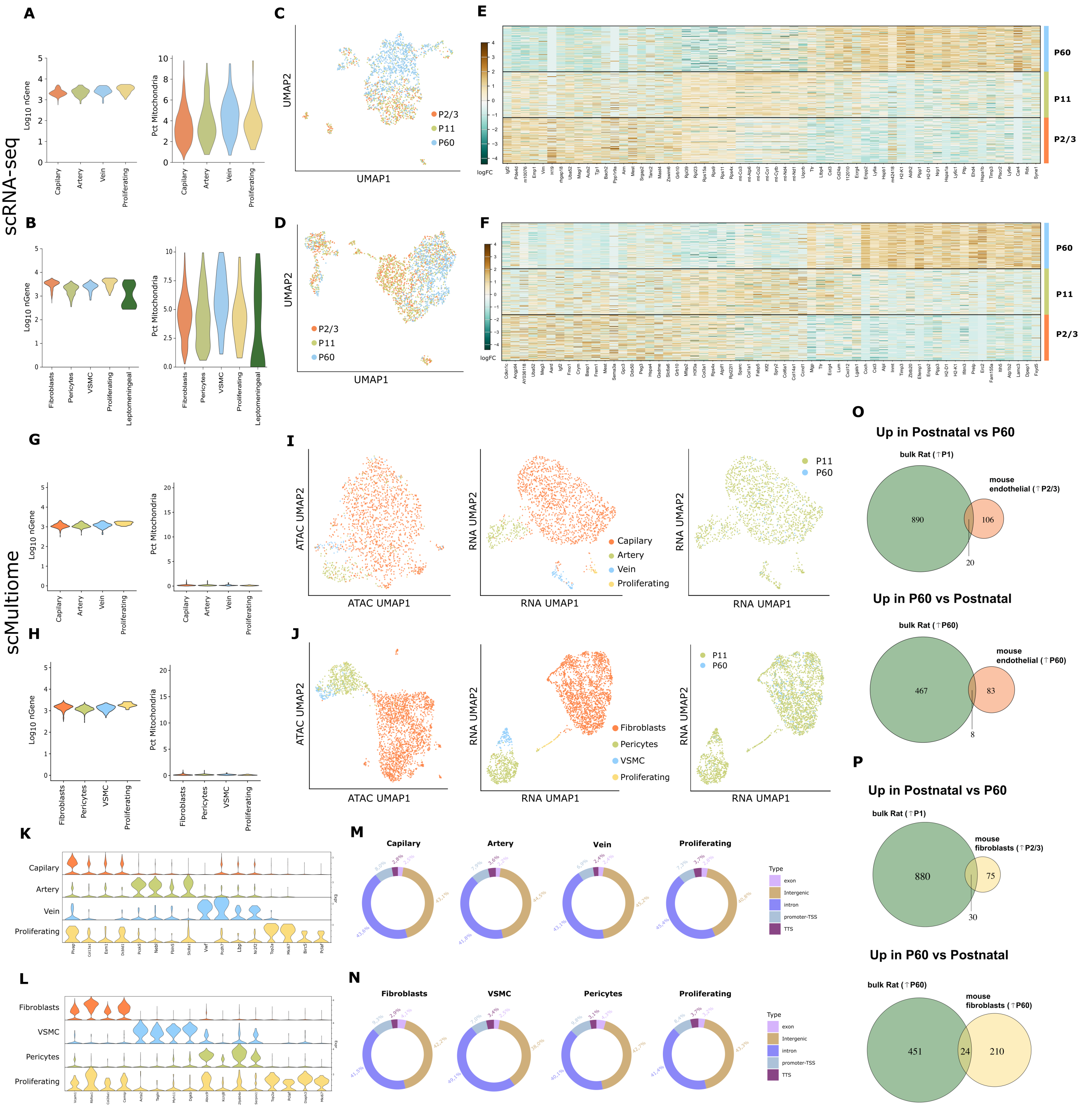


**Figure S4: scRNA-seq and scMultiome QC analysis for endothelial and mesenchymal cells, and differencially expression between postnatal and adults in different species.**

(A) and (B) Violin plots representing QC analysis of the endothelial and mesenchymal cells, respectively, in scRNA-seq, number of genes (log10nGene) and percentage (pct) of mithocondrial genes color-coded by cell-type.

(C )and (D) UMAP plots of endothelial and mesenchymal cells, respectively, analysed by scRNA-seq, color-coded by time point.

(E) and (F) Heatmap of differential expression genes in endothelial and mesenchymal cells, respectively, across different ages.

(G) and (H) Violin plots representing QC analysis of the endothelial and mesenchymal cells, respectively, in scMultiome, number of genes (log10nGene) and percentage (pct) of mithocondrial genes color-coded by cell-type.

(I) and (J) UMAP plots of endothelial and mesenchymal cells, respectively, analysed by scMultiome, for RNA and ATAC, color-coded by sample type and time point.

(K) and (L) Violin plot of median gene expression of four markers per cell population in the scMultiome data of endothelial and mesenchymal cells, respectively.

(M) and (N)  Distribution of the differentially accessible peaks amongst endothelial and mesenchymal cells, respectively.

(O)  and (O) Venn diagram representating the overlapp of genes up-regulated in postnatal stages versus adult P60 (and vice-versa) of both rat bulk RNA-seq and mouse endothelial and mesenchymal cells from scRNA-seq.

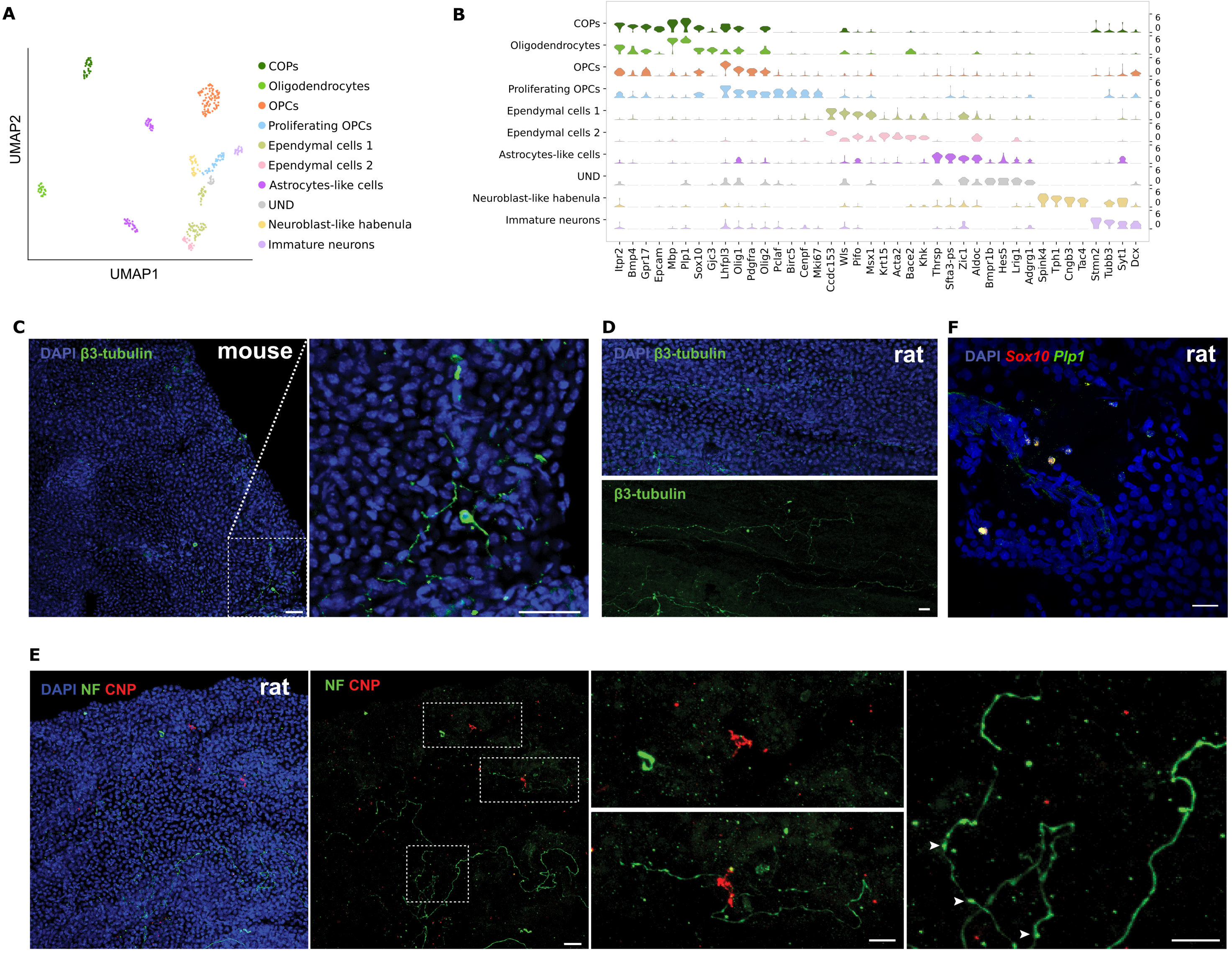


**Figure S5: Neuronal and glial cell types from rat and mouse lateral ventricle CP tissue**

(A) UMAP of neuronal and glial cell types color coded by cell type.

(B) Violin plots representing the median expression levels of markers for nine distinct populations.

(C) Immunofluorescence for β3-tubulin in adult mouse lateral ventricle CP. Representative images, n = 3 biologically independent adult mouse lateral ventricle CP. Scale bars, 20 µm.

(D) Immunofluorescence for β3-tubulin in adult rat lateral ventricle CP. Representative images, n = 3 biologically independent adult rat lateral ventricle CP. Scale bars, 20 µm.

(E). Immunofluorescence for CNP and NF in adult rat lateral ventricle CP. Dashed boxes shown at higher magnification highlight regions of interest. Representative images, n = 3 biologically independent adult rat lateral ventricle CP. Arrowheads point to axonal varicosities/boutons. Scale bars, 20 µm.

(F) RNAscope in situ hybridization in adult rat lateral ventricle CP with probes for *Plp1* and *Sox10*. Representative images, n = 3 biologically independent adult rat lateral ventricle CP. Scale bars, 20 µm.

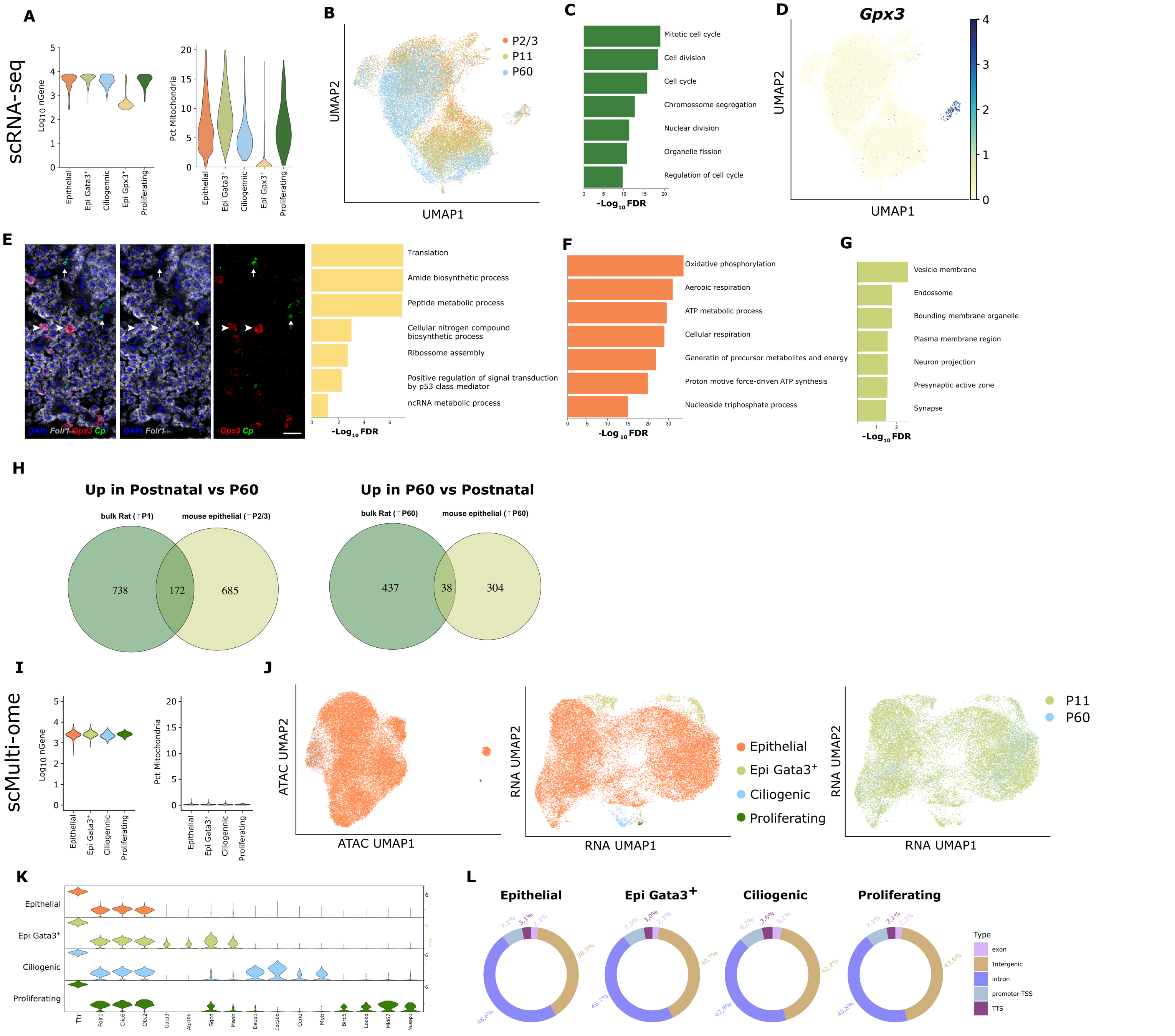


**Figure S6: scRNA-seq and scMulti-ome QC analysis for epithelial cells, gene ontology analysis for epithelial subtypes and overlapp of differencially expressed genes between postnatal and adults in different species.**

(A) Violin plots representing QC analysis of the five epithelial cell subtypes in scRNA-seq, number of genes (log10nGene), percentage (pct) of mithocondrial genes color-coded by cell-type.

(B) UMAP plots of epithelial cells analysed by scRNA-seq, color-coded by time point.

(C) GO analysis for the biological function on the top 50 genes enriched in proliferating (green) epithelial cells.

(D) UMAP plots depicting the expression of *Gpx3*.

(E) RNAscope ISH for *Folr1*$^+$*Cp*$^-$*Gpx3*$^+$ cells (arowheads) in whole mounts from of adult lateral ventricle CP and GO analysis for the biological function on the top 50 genes enriched in this subtype. Arrows represent *Folr1*$^+$*Gpx3*$^-$*Cp*$^+$ stromal cells. Representative images, n = 3 biologically independent adult mouse lateral ventricle CP. Scale bars, 20 μm.

(F) GO analysis for the biological function on the top 50 genes enriched in the main cluster (orange) of epithelial cells

(G) GO analysis for the cellular component on the top 50 genes enriched in the *Gata3*$^+$ epithelial subtype.

(H) Venn diagram representating the overlapp of genes up-regulated in postnatal stages versus adult P60 (and vice-versa) of both rat bulk RNA-seq and mouse epithelial cells from scRNA-seq.

(I) Violin plots representing QC analysis of the four epithelial cell populations in scMulti-ome, number of genes (log10nGene) and percentage (pct) of mithocondrial genes color-coded by cell-type.

(J) UMAP plots of epithelial cells analysed by scMulti-ome, for RNA and ATAC, color-coded by sample type and time point.

(K) Violin plot of median gene expression of four markers per cell subtype in the scMultiome data.

(L) Distribution of the differentially accessible of peaks amongst cell popupations.

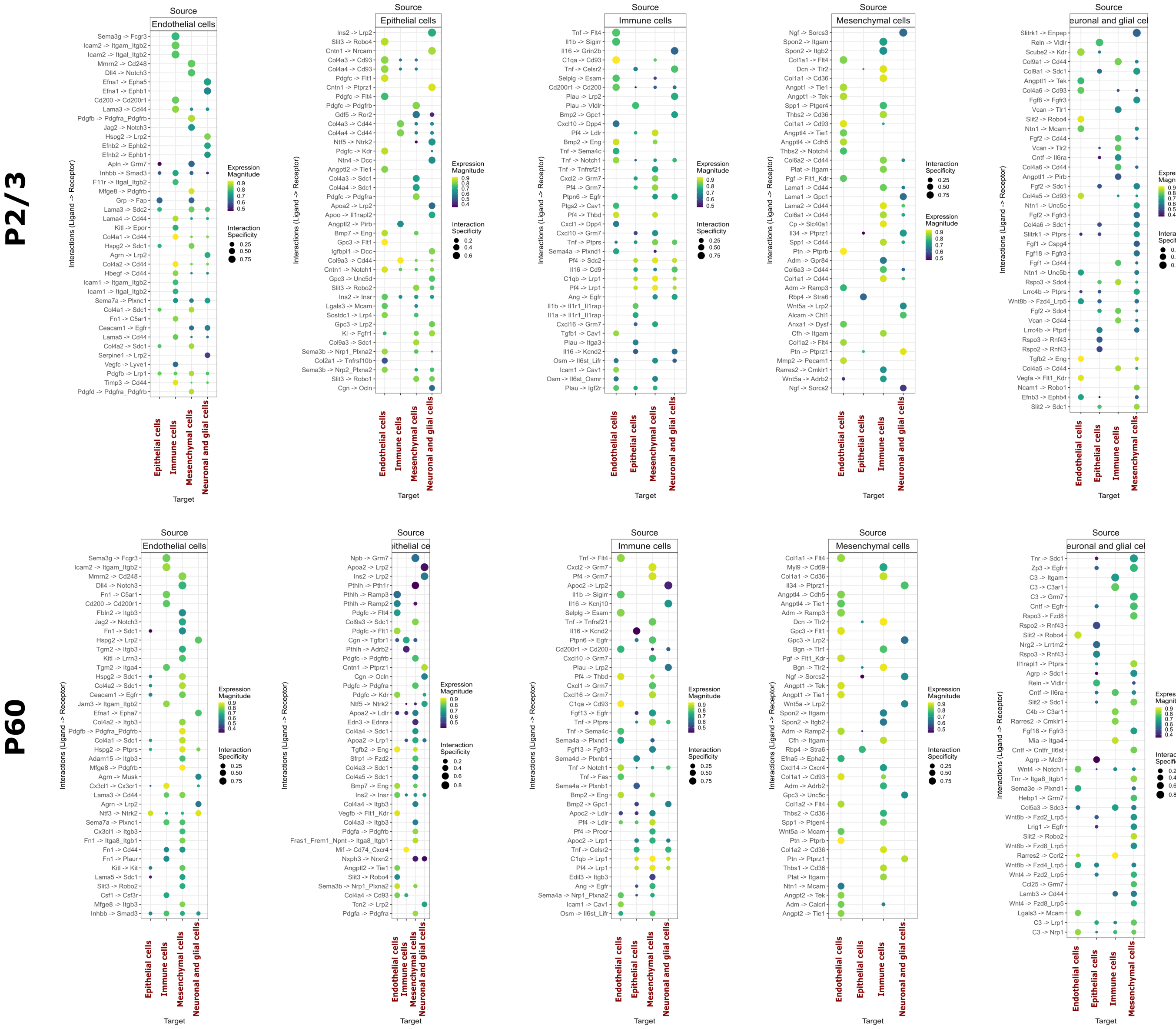


**Figure S7: Cell-cell communication inferred by LIANA, in the lateral ventricle CP cells from neonatal to adult stages.**
Dot plots for the 40 top interactions between the different cell types of the lateral ventricle choroid plexus in two different time points, P2/3 and P60. The magnitude of expression of the ligand and receptor, and the interaction specificity is represented on the right.
